# Inter-gastruloid heterogeneity revealed by single cell transcriptomics time course: implications for organoid based perturbation studies

**DOI:** 10.1101/2022.09.27.509783

**Authors:** Leah U. Rosen, L. Carine Stapel, Ricard Argelaguet, Charlie George Barker, Andrian Yang, Wolf Reik, John C. Marioni

**Affiliations:** European Molecular Biology Laboratory, European Bioinformatics Institute (EMBL-EBI), Hinxton, Cambridge, UK; Babraham Institute, Epigenetics Programme, Cambridge, UK; Altos Labs, Cambridge Institute of Science, Cambridge, UK; Wellcome Trust - Medical Research Council Institute of Metabolic Science, Metabolic Research Laboratories and Medical Research Council Metabolic Diseases Unit, University of Cambridge, Cambridge, UK; Wellcome Trust - Medical Research Council, Cambridge Stem Cell Institute, University of Cambridge, Cambridge, UK; Cancer Research UK Cambridge Institute, Li Ka Shing Centre, University of Cambridge, Cambridge, UK; Wellcome Sanger Institute, Wellcome Genome Campus, Hinxton, Cambridge, UK

## Abstract

Recent advances in organoid and genome editing technologies are allowing for perturbation experiments at an unprecedented scale. However, before doing such experiments it is important to understand the gene expression profile in each of the organoid’s cells, as well as how much heterogeneity there is between individual organoids. Here we characterise an organoid model of mouse gastrulation called gastruloids using single cell RNA-sequencing of individual organoids at half-day intervals between day 3 and day 5 of differentiation (roughly corresponding to E6.5-E8.75 *in vivo*). Our study reveals multiple differentiation trajectories that have hitherto not been characterised in gastruloids. Intriguingly, we observe that individual gastruloids displayed a strong bias towards producing either mesodermal (largely somitic) or ectodermal (specifically neural) cell types. This bifurcation is already seen at the earliest sampled time point, and is characterised by increased activity of WNT-associated pathways in mesodermally-biased gastruloids as compared to neurally-biased gastruloids. Notably, at day 5, mesodermal gastruloids show an increase in the proportion of neural cells, while neural gastruloids do not produce notably more mesodermal cells. This is in line with previous studies on how the balance between these cell types is regulated. We demonstrate using *in silico* simulations that without proper understanding of the inter-organoid heterogeneity, perturbation experiments have either very high false positive or negative rates, depending on the statistical model used. Thus in future studies, modelling of inter-organoid heterogeneity will be crucial when designing organoid-based perturbation studies.

**Highlights:**

- A single cell RNA-sequencing time course of day 3 to day 5 mouse gastruloids reveals multiple mesodermal and neural differentiation trajectories hitherto uncharacterised in gastruloids
- Single gastruloid, single cell RNA-sequencing of mouse gastruloids reveals that gastruloids are either mesodermally- or neurally-biased
- The two classes of gastruloid arise from differences in response strength to the WNT-agonist chiron
- At day 5, mesodermal gastruloids start making more neural cells, while neural gastruloids do not make more mesodermal cells, aligning with previously studied *in vivo* feedback loops
- We show using simulations that understanding interorganoid heterogeneity is a crucial consideration in the design and analysis of well-powered organoid-based perturbation studies

## Introduction

Organoids are a powerful tool for studying cell fate decisions in the context of development and disease. In particular, the ability to grow organoids in large quantities in combination with the ever expanding set of genetic editing tools makes them ideal for perturbation screens. However, before performing such perturbation screens it is important to not only understand how the organoid cell types relate to their *in vivo* counterparts, but also to quantify the inherent inter-organoid heterogeneity (Fleck et al., 2021).

A number of recent studies have characterised both gene expression and epigenetic states during mouse gastrulation at single cell resolution (Argelaguet et al., 2019; Guibentif et al., 2021; Mittnenzweig et al., 2021; Pijuan-Sala et al., 2019). This has yielded a large number of novel candidate genes and putative regulatory regions with potentially important roles in development. However, validating such a large number of candidates *in vivo* remains challenging, and instead organoid models are required. Gastruloids are an example of an organoid model that models mouse gastrulation (van den Brink et al., 2014). Each gastruloid is generated from a pool of 300 mouse embryonic stem cells that are aggregated for 48hrs, during which time they downregulate pluripotency factors (Baillie-Johnson et al., 2015). Subsequently, Wnt signalling is activated by a 24hr Chiron pulse, mimicking key signalling activity in the early mouse embryo (van den Brink et al., 2014). In response, gastruloids reproducibly break symmetry and produce all three main germ layers (ectoderm, mesoderm, endoderm) (van den Brink et al., 2014). Within five days of aggregation, gastruloids make posterior neural cells, neuromesodermal progenitors, and somitic tissues, as well as limited amounts of endoderm, PGC-like cells, cardiopharyngeal cells, and endothelium, but do not form anterior neural cells (Beccari et al., 2018; van den Brink et al., 2020). Extensions of the standard protocol exist that take gastruloids beyond the standard five days of differentiation (Beccari et al., 2018) or that bias their differentiation profile towards specific lineages, such as cardiac cell types (Rossi et al., 2021; Veenvliet et al., 2020).Thus far, however, neither the differentiation trajectories that lead to the generation of these cell types, nor intergastruloid heterogeneity have been profiled. Previous studies have either bulk sequenced pools of gastruloids (Beccari et al., 2018), profiled single cells from a pool of gastruloids (Anlaş et al., 2021; van den Brink et al., 2020; Rossi et al., 2021; Veenvliet et al., 2020) or spatially characterised very limited numbers of individual day five gastruloids (van den Brink et al., 2020).

More generally, the inherent variability of few organoid models has been rigorously studied. Quantifying inter-organoid heterogeneity is not only important for understanding the biology of the model, but also critical when using organoids in perturbation studies. Many previous studies comparing cell type abundance in perturbed versus wild type samples have assumed Poisson noise, an assumption that may well be violated (Haber et al., 2017; Ramachandran et al., 2019). Indeed, using single organoid sequencing, Gehling *et al*. reported that in hepatocyte organoids inter-organoid variability was very high relative to batch effects (Gehling et al., 2021). Similarly, using single cell RNA sequencing, Velasco *et al*. and Quadrato *et al*. showed that brain organoids had more than randomly expected variability (Quadrato et al., 2017; Velasco et al., 2019), though this variability was comparable to the one observed *in vivo* (Velasco et al., 2019). Other studies in epithelial intestinal organoids similarly showed a high degree of inter-organoid variability (Criss et al., 2021; Hof et al., 2021; Mohammadi et al., 2021). These findings suggest that there is significant, uncharacterised interorganoid heterogeneity across organoid systems, which is important to understand and subsequently model ahead of perturbation.

Here we profile day 3 to day 5 mouse gastruloids at half-day time points at single cell and single gastruloid resolution. We generated the first fine-grained time course of gastruloid development, identifying hitherto uncharacterised, earlier differentiation pathways. Furthermore, by assigning cells to individual gastruloids using MULTI-seq, we identified two classes of gastruloid with one making predominantly mesodermal and the other predominantly neural cell types, with further non-Poisson variance within each class, and identified potential drivers of this process. Finally, we demonstrate the importance of considering this heterogeneity when using organoids as a model for perturbation screens.

## Results

To investigate the developmental trajectory, and heterogeneity in gastruloid differentiation, we characterised single cells from individual gastruloids spanning differentiation day 3 to differentiation day 5 at half day time points by adapting MULTI-seq (McGinnis et al., 2019), an extension of the 10X single cell RNA-sequencing framework, for use with low cell numbers (Figure 1A). In MULTI-seq, cells from individual samples are barcoded prior to pooling, so that multiple samples can be sequenced together, and demultiplexed after sequencing (Figure 2A). 77,683 cells passed stringent quality control measures (Supplementary Figure S1A-H).

**Figure 1.**
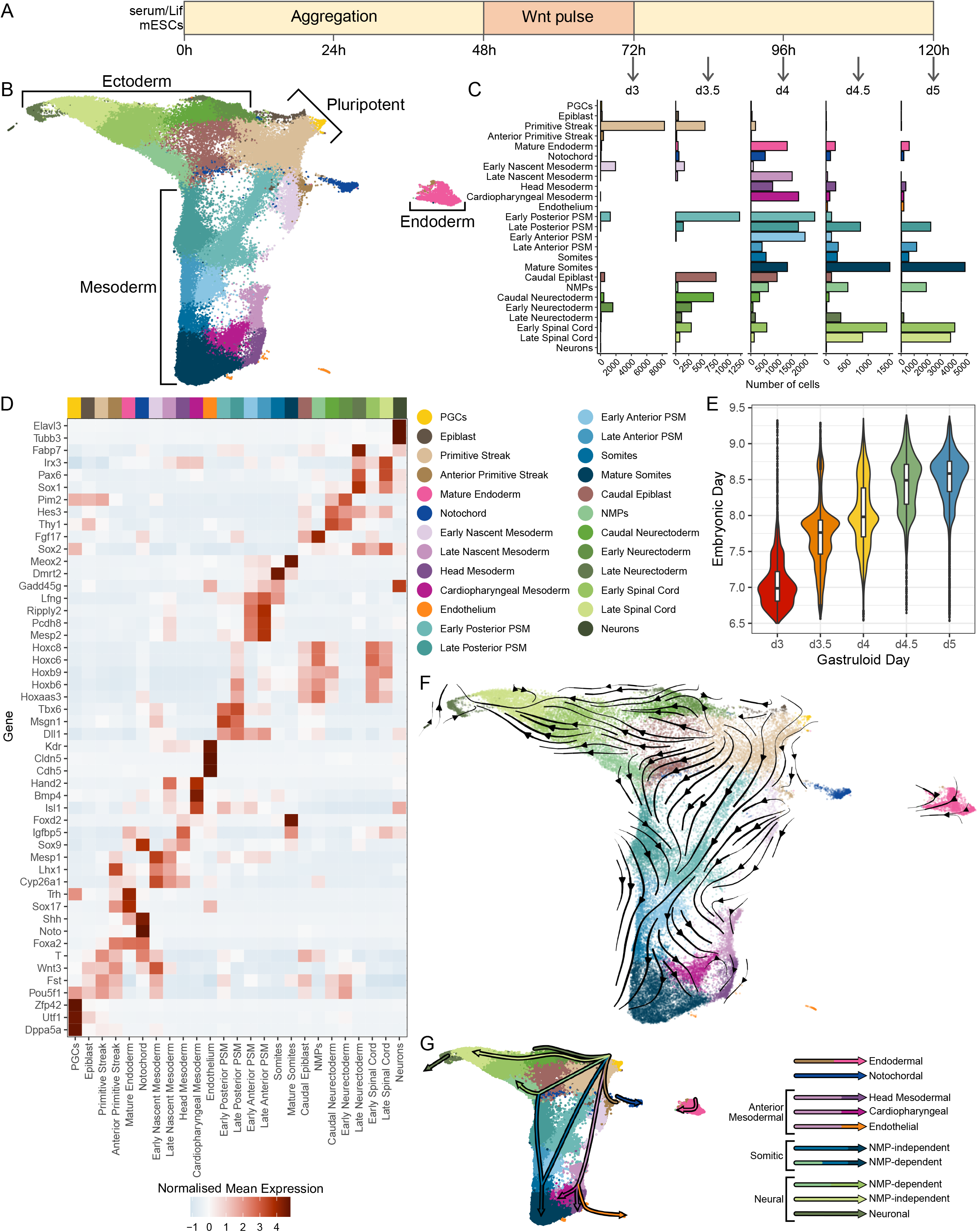
A single cell time course of mouse gastruloid development. A - Schematic of gastruloid development, along with indication of the five sampling time points. B - Uniform manifold approximation and projection (UMAP) plot showing all cells of the timecourse (77,683 cells). Cells are coloured by their annotated cell types, as explained in the legend below. C - Bar plot showing the number of cells assigned to each cell type at each of the five sampling time points. D - Heatmap showing standardised mean marker gene expression in each of the cell types. E - Violin and boxplot showing the embryonic day each gastruloid cell corresponds to, grouped by gastruloid sampling time point (day 3, n=15,144 cells, day 3.5, n=4,679 cells, day 4, n=17,718 cells, day 4.5, n=7,295 cells, day 5, n=20,811 cells). F - scVelo UMAP embedding streams overlaid on UMAP, coloured by assigned cell type, with a schematic of the 8 key trajectories. G - Schematic of the inferred trajectories on the UMAP.

**Figure 2.**
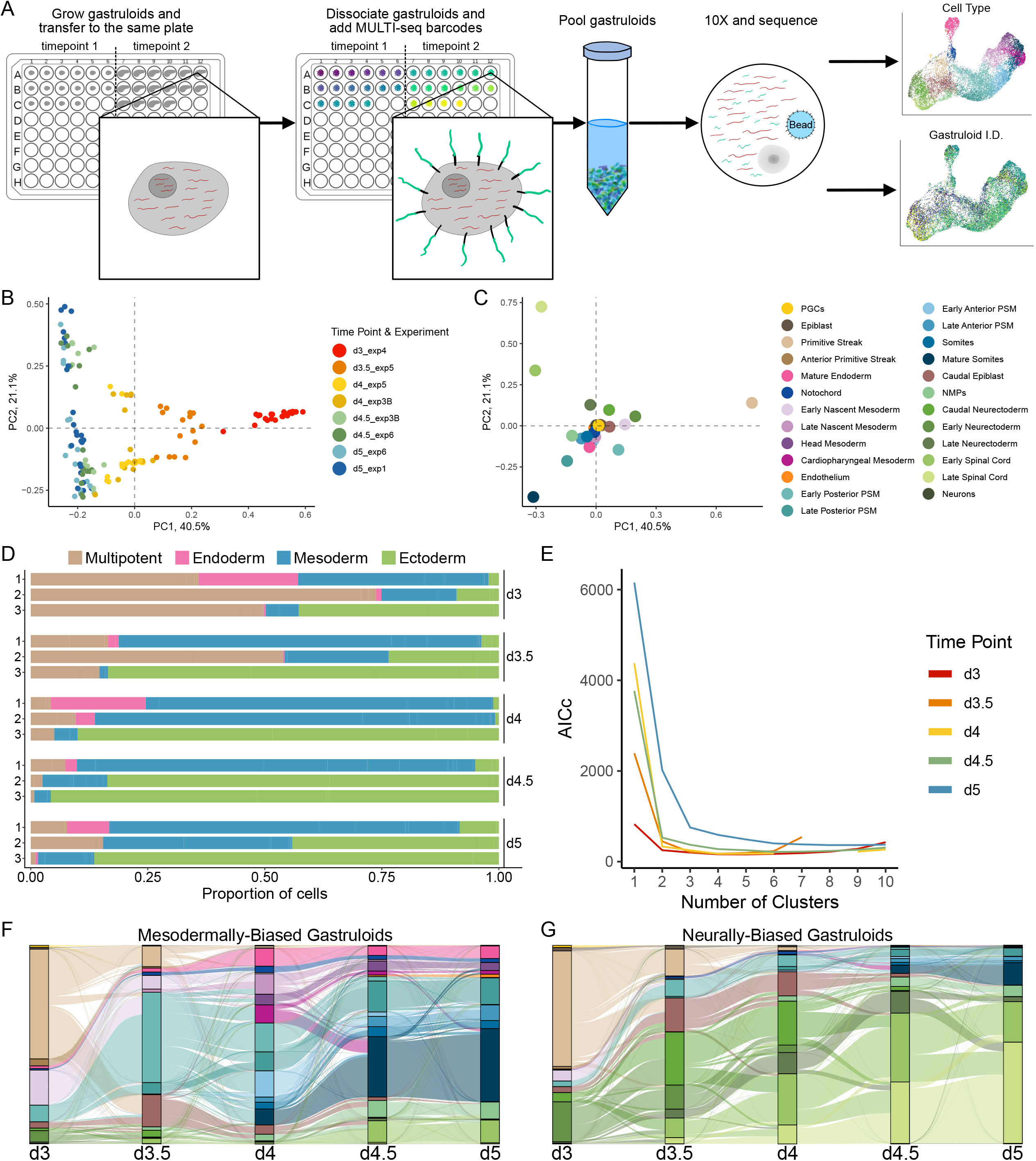
MULTI-seq reveals multiple classes of gastruloid. A - Overview of the gastruloid MULTI-seq experimental design. Two gastruloid time points are grown in two separate 96-well plates, started 12 hours apart. These are then replated into the same 96-well plate, where they are dissociated, and then labelled with MULTI-seq barcodes before being pooled and 10X sequenced. Barcodes and mRNA are separated based on size and sequenced separately (methods). After sequencing it is possible to know which gastruloid each cell came from. B - Embedding of individual gastruloids in a PCA space calculated based on the proportion of cells of each gastruloid assigned each cell type. Gastruloids are coloured based on the time point and experiment that they came from. C - Feature loadings of the PCA space shown in (B). Each point is a cell type, coloured according to the legend on the right. D - Bar plot of the proportion of cells of three representative gastruloids per time point that come from each lineage. Cell types are grouped into lineages for ease of visualisation. The selected gastruloids are highlighted in S5 and S6. E - Plot of the corrected Akaike Information Criterion (AICc) for each time point as increasing numbers of gastruloid clusters are fit. The AICc is based on a binomial distribution, with number of mesodermal versus number of neural cells tested. The lower the AICc, the more evidence for that number of clusters. F - Flow of cell types between time points according to Waddington OT inference in mesodermal gastruloids. G - Flow of cell types between time points according to Waddington OT inference in neural gastruloids.

### A time series of gastruloid development reveals multiple differentiation trajectories towards neural and mesodermal cell types

To identify the cell types that are present in gastruloids we generated a preliminary annotation by performing coarse-grained Louvain clustering combined with reference based cell type annotation using an atlas of mouse development spanning E6.5 to E9.5 (Imaz-Rosshandler et al., 2022). We further refined this annotation using marker genes. In total, we identified 25 cell types spanning primordial germ cells (PGCs), multipotent cell types, and other cell types derived from all three main germ layers (Figure 1B-D). Notably, gastruloid differentiation at all time points was highly reproducible between our replicates (each a pool of many gastruloids) (Supplementary Figure S1I) and gastruloid cell types at day 5 were concordant with previous single cell sequencing data (Supplementary Figure S2) (van den Brink et al., 2020), as well as qualitatively with Beccari *et al*. (Beccari et al., 2018), highlighting the reproducibility of the protocol. Consistent with previous observations in gastruloids and with *in vivo* data, few endodermal cells were observed relative to the number of mesodermal and ectodermal cells (Beccari et al., 2018; van den Brink et al., 2020). Furthermore, the ectodermal cells were primarily of a neural nature and the mesodermal cells followed either a somite or anterior mesoderm differentiation trajectory.

To characterise the differentiation dynamics of gastruloids in comparison to the mouse embryo we integrated our gastruloid time course data with the aforementioned mouse atlas. This revealed that day 3 gastruloids roughly corresponded to E6.75-E7.25, while day 3.5 gastruloids roughly corresponded to E7.5-E8.0 (Figure 1E). Day 4 gastruloids roughly corresponded to E7.75-E8.25, while both day 4.5 and day 5 gastruloids corresponded to E8.25-E8.75, with day 5 only slightly later. This is in agreement with data from bulk gastruloid and embryo sequencing (Beccari et al., 2018). However, the single cell data revealed that developmental time varied systematically by annotated gastruloid cell type (Supplementary Figure S3A). For example, spinal cord cells can be found in gastruloids as early as day 3, whereas this cell type is only present from E8.25 in mouse embryos.

Leveraging the time course data, we inferred differentiation trajectories using CellRank (Lange et al., 2022), which integrates RNA Velocity (Bergen et al., 2020; La Manno et al., 2018) and connectivity graph inferences (Figure 1F). This demonstrated that gastruloids followed three separate mesoderm differentiation trajectories: An anterior mesodermal trajectory that gives rise to head mesoderm, cardiopharyngeal mesoderm, and endothelium; a neuromesodermal progenitor (NMP)-dependent somitic trajectory, and an NMP-independent somitic trajectory (Figure 1G). Furthermore, it suggests that there was an NMP-independent neural pathway giving rise to non-spinal cord neurons as well as spinal cord, together with an NMP-dependent pathway giving rise to spinal cord (Figure 1G). These inferences are in line with previously known lineage relationships.

Thus, our single cell analysis of a time course of gastruloids has revealed the differentiation trajectories that are followed in gastruloids in detail, highlighting new avenues for the use of gastruloids to model mouse development. Our data also demonstrate heterochronicity in gastruloid cell type differentiation compared to equivalent *in vivo* data.

### Single gastruloid sequencing reveals two classes of gastruloid

To investigate the heterogeneity between gastruloids, we next assigned cells to individual gastruloids using their MULTI-seq barcode identity (Figure 2A). In total, we assigned cells to 136 individual gastruloids across our entire time series (d3, d3.5, d4, d4.5, d5; Supplementary Figure S4A) with a median of between 182 and 473 cells per gastruloid being recovered (Supplementary Figure S4B).

Upon first inspection, it was immediately clear that substantial heterogeneity in cell type proportions exists between gastruloids (Supplementary Figure S5). Therefore, we calculated the individual cell type proportions for each gastruloid, and plotted these in PCA space (Figure 2B,C, Supplementary Figure S6P). The PCA embedding not only separated the gastruloids according to their respective differentiation days (along the first principal component), but also revealed that gastruloids could be split into two classes along the second principal component, one making predominantly mesodermal, and the other making predominantly neural cell types. This is in line with Beccari *et al*. not detecting expression of mesodermal and endodermal markers in all profiled gastruloids using in situ hybridisation (Beccari et al., 2018).

Performing PCA on only differentiation day 3 (24hr post chiron pulse) gastruloids revealed, even at this earliest time point, a clear bifurcation into mesodermal and neural gastruloids (Supplementary Figure S5A, S6A, F, K), with even clearer separations being observed at later time points (Supplementary Figure S6A-O). Interestingly, despite these substantial differences, all gastruloid classes contained cells from all three main germ layers, just at vastly different proportions (Figure 2D, Supplementary Figures S5–7). We confirmed the statistical significance of these two classes by subsetting the gastruloids to only cells assigned a mesodermal or neural identity (Supplementary Figure S6Q), and running k-means on the proportions with k between 1 and 10 clusters. We then calculated the corrected Akaike Information Criterion (AICc) for this number of clusters, based on a binomial distribution on the number of mesodermal and neural cells per gastruloid (Figure 2E). This confirmed the bifurcation, with clear evidence for two clusters of gastruloids being preferred over one cluster at all time points. Interestingly, there was also clear evidence that at differentiation day 5 there were three clusters instead of two clusters (Figure 2E). Taken together with the PCA analysis (Supplementary Figure 6J,P), this third class of gastruloids appears to be an intermediate between the mesodermally-biased and neurally-biased classes (Supplementary Figures 6J and 6P). We thus classified each gastruloid as either being mesodermal or neural, or, in the case of the third day 5 cluster, “intermediate” (Supplementary Figure S6F-I).

We then performed Waddington OT trajectory inference as implemented in CellRank (Lange et al., 2022; Schiebinger et al., 2019) in mesodermal and neural gastruloids separately, excluding the day 5 intermediate gastruloids owing to their ambiguous day 4.5 origin, to investigate transitions between subsequent time points (Figure 2F,G), as well as an integrated CellRank analysis of RNA velocity, connectivity, and Waddington OT (Supplementary Figure S8A,B). This confirmed the velocity-based CellRank findings (Figure 1F), but interestingly identified NMPs as a terminal state (Supplementary Figure S8C,D), with the somites and spinal cord predominantly derived from the NMP-independent trajectories. Furthermore, Waddington OT inferred that at day 4 in mesodermal gastruloids, a small pool of caudal epiblast, NMPs and spinal cord cells proliferate and differentiate, driving the convergence towards a more balanced cell type distribution at later time points. Similarly, in neural gastruloids a population of early posterior presomitic mesoderm cells persists between day 3.5 and day 4, which subsequently differentiate, though with less relative proliferation (Figure 2F,G, Supplementary Figure S8C,D).

### Varying WNT responses at day 3 explain gastruloid bifurcation

To investigate the origins of the intergastruloid heterogeneity, we used Weighted Gene Co-Expression Network Analysis (WGCNA) to find biological properties significantly associated with our phenotype of interesty (Barker et al., 2022; Langfelder and Horvath, 2008). For this we first ran single cell WGCNA (Feregrino and Tschopp, 2021) on only the day 3 MULTI-seq cells. This revealed 34 gene coexpression modules, with each module containing a mutually exclusive set of between 17 and 499 genes. We then pseudobulked the day 3 gastruloids and calculated how much of the variance in the pseudobulked gastruloid by gene matrix each module explained. Based on this, we identified 20 modules that explained a significant amount of variance (p<0.05 after Bonferroni correction). These modules could be clearly separated into those that were enriched in either neural or mesodermal gastruloids, with three mesodermal and three neural modules being particularly associated with gastruloids near the start of the bifurcation (Figure 3A). We classified these as “early mesodermal” and “early neural” modules, and combined the genes in each of the two groups. Comparative GO enrichment analysis of the two groups found five significant terms (Fisher exact test on GO terms that were significant at an adjusted p-value threshold of 0.05 in at least one of the two classes, with a p-value threshold of 0.05 and an FDR threshold of 0.1; Figure 3B). These terms were all enriched in the mesodermal modules and not in the neural modules, and are associated with Wnt signalling and establishment of planar cell polarity. WNT signalling is the key pathway of gastrulation, and AP-axis establishment an important consequence thereof, which suggests that the mesodermal gastruloids receive new signals that stimulate gastrulation. Given that gastruloids are generated using a chiron pulse (a Wnt agonist) between differentiation days 2 and 3, the higher WNT signalling in day 3 mesodermally-biased gastruloids suggests that these gastruloids responded more strongly to the chiron pulse, while gastruloids that respond less strongly to these signals defaulted along a more neural trajectory. This is in line with previous, primarily *in vitro*, studies that find that the neural fate is the default (Tropepe et al., 2001). Interestingly, neuron differentiation is also associated with mesodermal gastruloids. However, this is not unexpected, since WNT signalling promotes neural progenitor maintenance and neuron maturation (Edri et al., 2019). This could either be caused by differences in chiron pulse strength, or by differences in competence to respond to chiron signals. This interpretation agrees with the findings of Sáez *et al*. that in a 2D differentiation system, small differences in chiron concentration can lead to the formation of either predominantly posterior neural or predominantly mesodermal tissues (Sáez et al., 2022).

**Figure 3.**
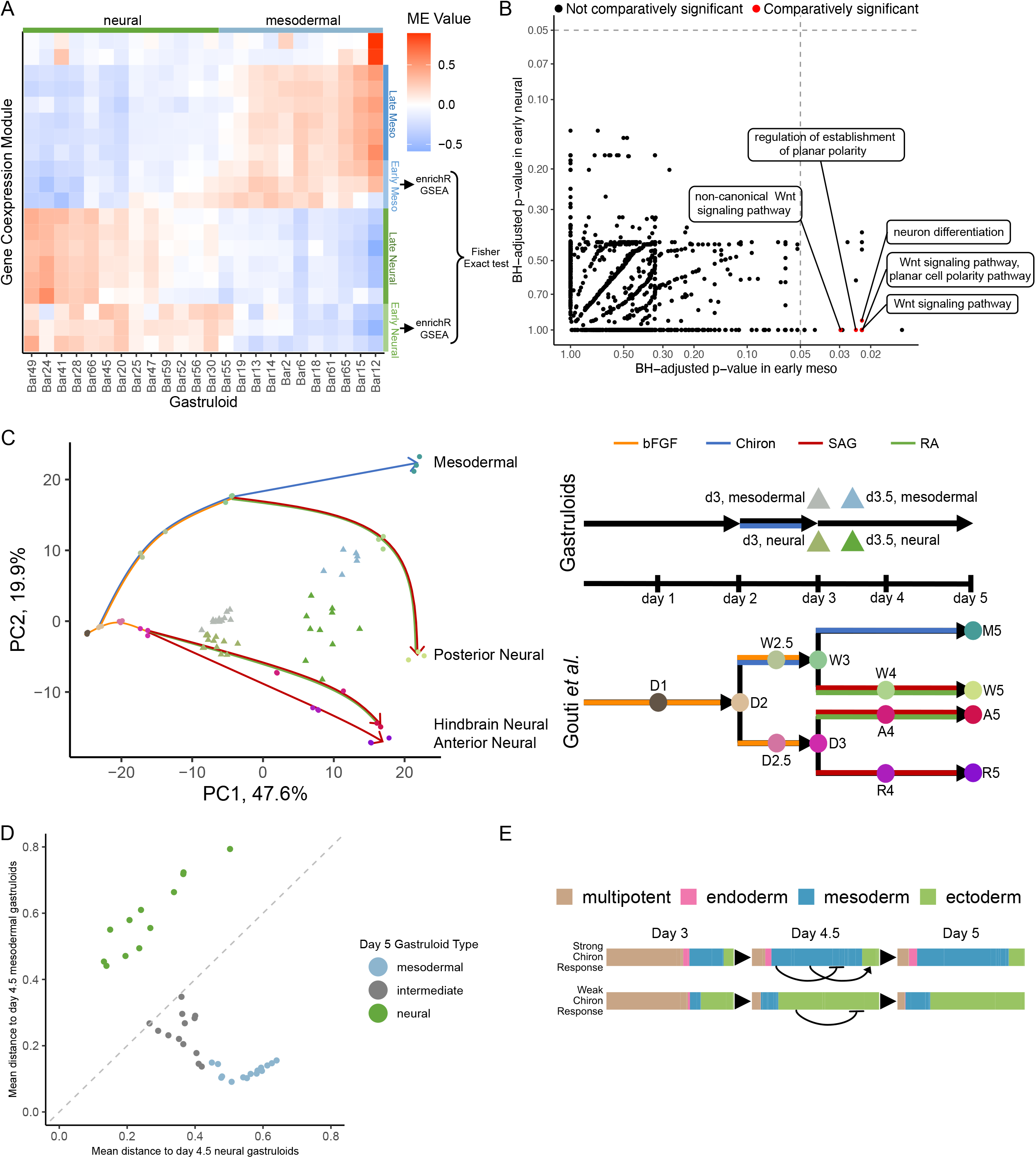
Separate trajectory inference for each gastruloid class. A - Heatmap showing the value of each module eigengene in the pseudobulked day 3 gastruloids. scWGCNA was run on the single cell gastruloid data, and modules that explained significant amounts of variance between gastruloids selected. Gastruloids were sorted based on their ordering along the principal curve in the day 3 cell type proportion PCA space, and their class annotated. Modules are sorted based on hierarchical clustering of the gastruloid by module eigengene matrix. Modules were manually classified. B - Plot showing all GO terms, and their Benjamini-Hochberg adjusted p-value based on enrichR GO enrichment analysis in the early mesodermal and early neural modules. GO terms are coloured based on whether they were differentially significant in the early mesodermal versus the early neural modules based on a Fisher exact test. The significant GO terms are labelled. C - Limma-integrated gene expression PCA space of Gouti *et al*. bulk RNA-sequencing data of 2D differentiated cells, and individual pseudobulked day 3 and day 3.5 gastruloids. Gouti *et al*. samples are circles, coloured based on their time point and differentiation conditions. Individual gastruloids are triangles, coloured based on their time point and whether they were assigned a mesodermal or neural identity. Arrows show the Gouti *et al*. differentiation conditions. D - Plot of individual day 5 gastruloids showing each gastruloids mean distance, in cell type proportion PCA space, to the day 4.5 mesodermal gastruloids, and the day 4.5 neural gastruloids. Gastruloids are coloured based on their assigned class, and the x=y line is highlighted. E - Schematic of the proposed model explaining the increased proliferation of neural cells in mesodermal gastruloids at later differentiation time points.

To further investigate the hypothesis that differences in chiron response are the cause of this bifurcation we compared our gastruloids to bulk RNA-seq data where cells in similar culture conditions were either differentiated towards anterior neural fates in the absence of chiron, or towards mesodermal and posterior neural fates in the presence of chiron (Gouti et al., 2014). For this we pseudobulked each day 3 and day 3.5 gastruloid and then projected it into the bulk RNA-seq PCA space (Figure 3C). We found that day 3 neural gastruloids fell along the trajectory of cells cultured without chiron between day 3 and day 4. In contrast, day 3 mesodermal gastruloids were more similar to the day 3 cells cultured with chiron. By day 3.5 all gastruloids spanned a range between the day 4 non-chiron cultured cells and day 4 cells that had been cultured with chiron between day 2 and day 3 (the condition most similar to gastruloids), with the mesodermal gastruloids still more similar to those that had been cultured with chiron, and the neural gastruloids more similar to those cultured without. This provides additional evidence that the difference between mesodermal and neural gastruloids is driven by differing strengths of the chiron response.

### Day 5 intermediate gastruloids are derived from mesodermally-biased gastruloids

To better understand the previously-identified day 5 intermediate gastruloid class, we ran PCA on the cell type proportions of all day 4.5 and day 5 gastruloids. Interestingly, all intermediate gastruloids are closer in PCA space to the day 4.5 mesodermal gastruloids than they are to the day 4.5 neural gastruloids (Figure 3D). This means that there would have to be less change in the cell type proportions if the intermediate gastruloids were derived exclusively from the mesodermal lineage, than if they were derived from both mesodermally- and neurally-biased gastruloids. Additionally, when comparing each day 5 gastruloid’s mean distance to the day 4.5 neural versus the day 4.5 mesodermal gastruloids, the intermediate gastruloids form a continuous trajectory with the day 5 mesodermal gastruloids, while the day 5 neural gastruloids cluster separately (Figure 3D). Combined with the fact that the intermediate gastruloids are further from the day 4.5 mesodermal gastruloids than are the day 5 mesodermal gastruloids (Figure 3D), this suggests that the intermediate gastruloids are developmentally later forms of the day 5 mesodermal gastruloids. Furthermore, as the day 5 neural gastruloids develop further away from the day 4.5 neural gastruloids, they also move further in PCA space from the day 4.5 mesodermal gastruloids (Figure 3D), suggesting that they acquire a stronger neural identity. Together, this suggests that at the latest developmental time points, mesodermal gastruloids begin to produce more neural cells, while neural gastruloids do not produce notably more mesodermal cells.

These observations are consistent with a previous study that investigated the gene regulatory networks that underpin the choice between posterior mesoderm and neural differentiation and proposed that an abundance of mesodermal cells stimulates neural differentiation through the induction of retinoic acid signalling (Figure 3E) (Gouti et al., 2017). Conversely, though a low abundance of mesodermal cells reduces retinoic acid signalling and neural cell differentiation, it does not stimulate Wnt signalling and mesodermal differentiation. Thus, neural gastruloids which display low levels of Wnt signalling are not able to increase mesoderm differentiation and remain mostly neural. Importantly, both gastruloid classes make all three main germ layers, thus there are small numbers of neural cells in mesodermal gastruloids that are able to differentiate, as shown by the Waddington OT analysis (Figure 2F). Similarly there are small numbers of mesodermal cells in the neural gastruloids that are able to proliferate (Figure 2G). Consistent with the proposed explanation, mesodermal cells in neural gastruloids proliferate less relative to the neural cells do in neural gastruloids than the neural cells in mesodermal gastruloids do, relative to the mesodermal cells in mesodermal gastruloids (Supplementary Figure 8A,B).

### Interorganoid heterogeneity critically influences statistical power in perturbation experiments

Increasingly, organoids are being used as models in which to perform perturbation experiments with single cell readout, due to the relative ease of perturbing them, combined with the relative ease of generating large numbers of organoids. However, thus far the interorganoid heterogeneity has not been considered in these experiments beyond generating standard wild type controls, this is particularly pertinent in a diverse, differentiating, 3D system like the gastruloid. In its simplest form, an organoid perturbation experiment with single cell readout involves generating some number of wild type samples, and some number of samples from a perturbed condition. Each sample consists of a pool of organoids, that are sequenced using single cell RNA sequencing, cell types assigned, and differential abundance testing, usually using a Poisson linear model, is performed (Figure 4A).

**Figure 4.**
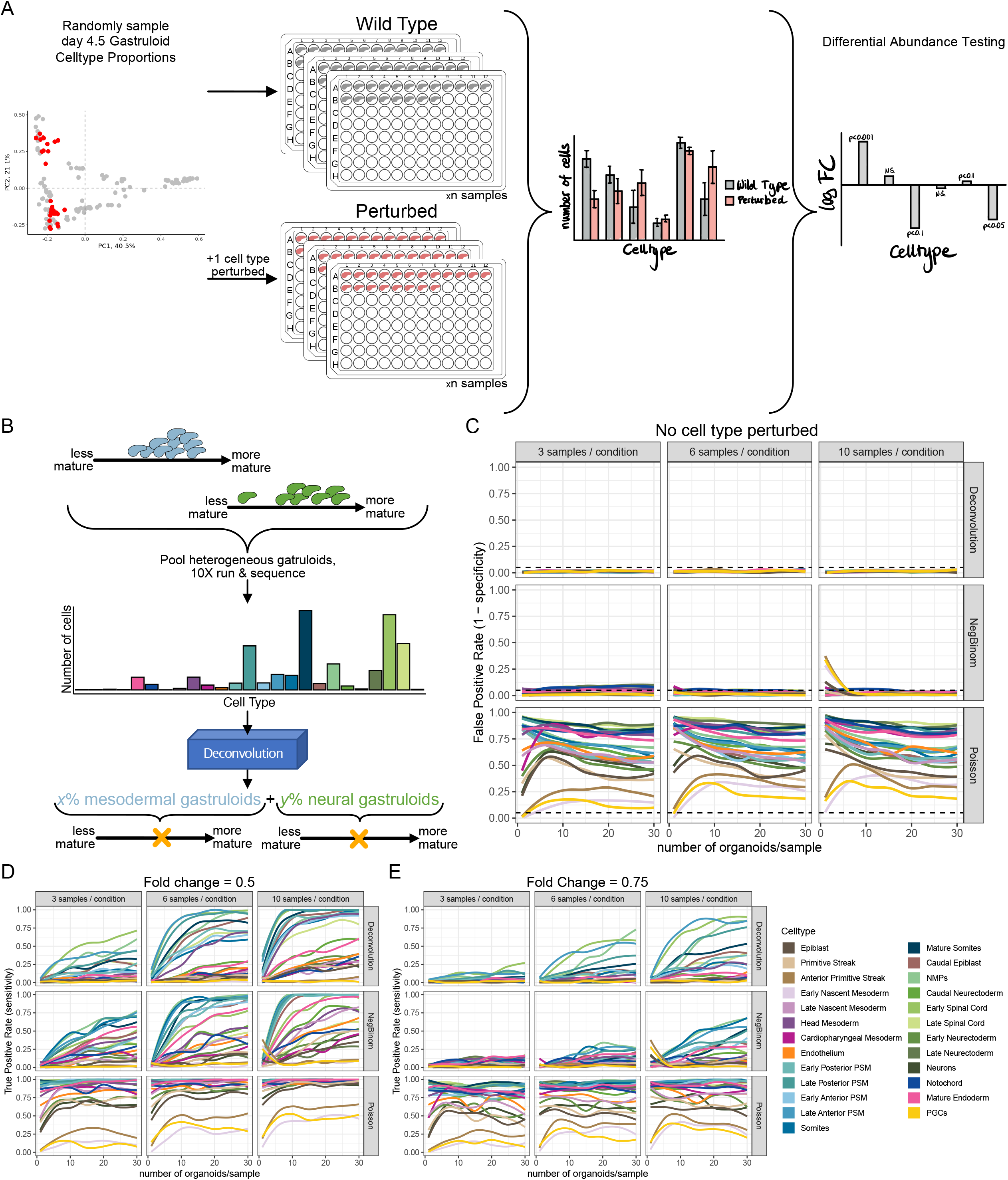
Effect of inter-organoid variability on statistical power in perturbation experiments. A - Schematic of the simulation and DA testing approach. G gastruloids are simulated twice the number of sample times, by randomly sampling the cell type proportion of day 4.5 gastruloids with replacement. In the perturbed condition, one cell type is downregulated at some fold change, and then the entire proportion matrix re-normalised to sum to 1. Based on the number of cells per cell type in each sample, either a Poisson, a negative binomial, or a custom deconvolution-based differential abundance test is performed. B - Schematic of the deconvolution-based differential abundance testing method. The input distribution is modelled from the observed, aggregate cell counts, by modelling the proportion of cells coming from mesodermal gastruloids, and the proportion coming from neural gastruloids, as well as how mature the mesodermal and neural gastruloids are, respectively. C - False positive rate of the three differential abundance testing approaches for varying numbers of samples per condition, and gastruloids per sample, when no cell type is perturbed. Colours indicate which cell type is being tested, according to the legend in the bottom right. The horizontal line shows the position of a 0.05 false positive rate. The rate is based on 100 simulations. D - True positive rate of the three different differential abundance testing approaches for detecting that a given cell type is differentially abundant at a fold change of 0.5, for varying numbers of samples per condition and gastruloids per sample. Colours indicate which cell type is differentially abundant, according to the legend to the right. The rate is based on 100 simulations. E - As E, but the given cell type is differentially abundant at a fold change of 0.75.

When designing these studies, there are a number of key parameter choices, including the number of samples and the number of organoids per sample. These choices are critically influenced by the expected signal to noise ratio, i.e., the strength of a perturbation effect relative to the inherent null distribution. To demonstrate this using the example of gastruloids, we simulated such a perturbation experiment with varying parameter choices. To this end, we bootstrap sampled cell type proportions from day 4.5 gastruloids that were pooled *in silico* to form n WT samples and n perturbed samples. In the perturbed samples a single cell type was decreased in abundance at various fold changes.

Using a Poisson model for differential abundance testing leads to many false positives (Figure 4C), while using a negative binomial model for differential abundance testing leads to many false negatives (Figure 4D,E), across a typical range of sample and gastruloid number choices when the fold change is 0.5, and across a wide range when the fold change is 0.75. However, given a large enough number of gastruloids per sample and large enough number of samples, the negative binomial model is able to gain power to detect fold changes of 0.5. This shows that knowing the inherent variability that is to be expected, as well as the expected fold change, is crucial before designing a perturbation experiment.

However, it is possible to improve the sensitivity by leveraging prior knowledge about the cell type composition of individual gastruloids. Here we propose a model inspired by cell type deconvolution (Young et al., 2021), in which we explicitly model the structure of the inter-gastruloid heterogeneity. Each sample in the perturbation experiment is comprised of multiple gastruloids (Figure 4A). These gastruloids are either mesodermally- or neurally-biased. In addition there is heterogeneity within each class largely driven by how mature each individual gastruloid is. In the proposed model, we deconvolve what proportion of the cells of each sample come from each gastruloid class, and within each class, we model the continuous variance (Figure 4B). We then use this inferred distribution to find cell types that diverge from the expected heterogeneity structure in the perturbed samples relative to the wild type samples, which are called as differentially abundant. This model had a lower false positive rate when no cell type was perturbed than both the Poisson and negative binomial models (Figure 4C), while being more sensitive to small changes in cell type abundance than the negative binomial model (Figure 4E). While this model does have systematic false discoveries, caused by attempting to fit mean values, when one cell type is perturbed, these false discoveries can easily be identified and removed (Supplementary Figure 9A). Therefore, understanding the underlying heterogeneity of the organoid model allows for both powerful experimental design, as well as more power in the downstream statistical analysis.

## Discussion

Our study provides the first single cell time course of gastruloid development, revealing hitherto unknown gatruloid differentiation trajectories. In addition, our results show that gastruloids develop along one of two trajectories, either making primarily mesodermal, or primarily neural cell types, with mesodermal gastruloids starting to make more neural cells at day 5. This interorganoid heterogeneity not only reveals key features of gastruloid biology, but the convergence of day 5 mesodermal, but not neural, gastruloids makes them an attractive system in which to study the mechanisms by which embryos make the correct proportion of each cell type.

Organoids are increasingly used as systems in which to perform perturbation screens. Our findings demonstrate that it is critical to quantify the inherent interorganoid heterogeneity before performing such perturbation screens, or else the variance will be incorrectly estimated leading to a large number of false positives or false negatives, depending on the statistical model used. However, we show that the structure of the heterogeneity can be leveraged using a deconvolution-based differential abundance testing approach to obtain higher power. Importantly, considering inter-organoid heterogeneity is important beyond differential abundance testing. Firstly, it is important to also consider whether a perturbation may have different effects in different classes of organoid. Secondly, if a perturbation makes only one group of organoid less fit, this could lead to unaffected cell types being assumed to be affected.

More broadly, organoid models are being used to study, via large-scale CRISPR screens, the mechanisms that underpin disease, including in the context of personalised medicine. Our study provides a two-pronged framework for performing such screens: one part focuses on understanding the intrinsic heterogeneity of the system, with the second involving a larger CRISPR screen involving pooling of individual organoids into a single sample. Critically, our results demonstrate that data generated by the screen can only be properly analysed using the first component, outlining a path forward for extracting the most biological insight from such studies.

## Code and Data availability

All code is available at https://github.com/MarioniLab/gastruloids2022 and all RNA and MULTI sequencing datasets are deposited in the Gene Expression Omnibus (GEO) under accession number GSE212050. The data can be interactively explored at https://marionilab.cruk.cam.ac.uk/GastruloidTimecourseShiny/.

## Acknowledgements

We thank Nicole Forrester and Paula Kokko-Gonzales of the Babraham Next Generation Sequencing Facility for 10X sequencing, Katarzyna Kania and Paul Coupland from the CRUK-CI sequencing facility for 10X sequencing, and Felix Krueger of the Babraham Bioinformatics Group for processing and managing sequencing data. Christopher McGinnis for sharing MULTI-seq reagents and advice on performing MULTI-seq with low cell input. We would also like to thank Alfonso Martinez Arias, James Briscoe, Naomi Morris, and Valerie Wilson for discussions on gastruloid biology, the cell types they make, and the signalling pathways involved in the two gastruloid classes. We thank Peter Peneder, Ioannis Kafetzopoulos, and Alfonso Martinez Arias for their thoughtful comments on the manuscript. We would further like to thank Ivan Imaz-Rosshandler for sharing the processed extended mouse atlas data with us, Alice Santambrogio and Ioannis Kafetzopoulos for discussions, Shila Ghazanfar for advice on comparing the early mesodermal to early neural WGCNA modules, Emma Dann for advice on differential abundance testing, and for sharing her Poisson and negative binomial differential abundance testing code, Maria Kalyva for discussions on cell type deconvolution, and for help with understanding the Youg *et al*. method, Leah McHugh for discussions on how to present the results, Rebecca Berrens for discussions about PGCs, Daniel Keitley for discussions about how to compare *in vivo* to *in vitro* cells, and Artem Lomakin for discussions about how to model gastruloid cell type proportions. LUR was funded by the EMBL International PhD Programme, and is a member of Darwin College of the University of Cambridge. LCS was supported by a Marie Sklodowska-Curie Individual Fellowship (EpiNoise 798499) funded by the European Commission under the H2020 Programme and was a Junior Research Fellow at Wolfson College of the University of Cambridge. This work was funded as part of a Wellcome Trust Grant (220379/Z/20/Z) awarded to W.R., J.C.M., and B. Göttgens; and as part of an ERC grant (EpiCell lineage 883798) funded under the H2020 Programme awarded to W.R.

## Author Contributions

LCS and WR conceived the study. LCS performed the gastruloid differentiation and single cell sequencing experiments. LUR, RA, and AY analysed and visualised the data. CGB provided code for and advice on the computational analysis of the day 3 gastruloid signalling pathways. LUR, LCS, WR, and JCM interpreted results and drafted the manuscript. WR and JCM supervised the project. All authors read and approved the final manuscript.

## Methods

### Mouse ESC and gastruloid culture

Mouse E14Tg2A (E14) Embroynic Stem Cells (ESCs) obtained from the Cambridge Stem Cell Institute (CSCI) were cultured in ESLIF medium (DMEM (Life Technologies, 10566-016), 15% fetal bovine serum (Sigma-Aldrich, Lot #BCCC3714), 1x MEM Non-Essential Amino Acids (Life Technologies, 11140050), 2 mM GlutaMAX (Life Technologies, 35050061), 10 U/ml Penicillin-Streptomycin (Life Technologies, 15140122), 0.1 mM 2-Mercaptoethanol (Life Technologies, 31350010), and 10 ng/ml mLIF (Department of Biochemistry, University of Cambridge)) on gelatin-coated (0.1% in H2O) tissue culture treated 10 cm plates in a humidified incubator at 37 °C, 5% CO2. Cells were passaged every other day.

Gastruloids were grown as described previously (Girgin *et al*. 2018 Protocol Exchange). Briefly, ESLIF-grown mouse ESCs were dissociated to a single cell suspension using Trypsin-EDTA (Thermo Fisher Scientific, 25300096). Trypsin was inactivated in 5 ml ESLIF medium and cells were spun down. The cell pellet was washed and spun down in 5 ml pre-warmed PBS (14190144) twice to remove remaining traces of ESLIF medium. After the second wash, cells were resuspended in 5 ml N2B27 medium (50/50 mix of DMEM-F12 (Thermo Fisher Scientific, 11320033) and Neurobasal (Thermo Fisher Scientific, 21103049), supplemented with 0.5x N-2 Supplement (Cell Therapy Systems, A1370701), 0.5x B-27 (Thermo Fisher Scientific, 12587010), 2 mM GlutaMAX, 10 U/ml Penicillin-Streptomycin and 0.1 mM 2-Mercaptoethanol) and counted using a haemocytometer. Cells were diluted in N2B27 medium to a density of 7500 cells/ml. This cell suspension was transferred to a reservoir and 40 μl of the suspension was added to each well of a U-bottom suspension culture 96-well plate (Greiner Bio-One, 650185) to reach a cell density of 300 cells/well. These cells were left to aggregate for 48 hours, at which point 150 μl of N2B27 medium with 3 μM of the GSK3 inhibitor CHIR99021 (Chiron, Department of Biochemistry, University of Cambridge) was added. Subsequently, 150 μl of the medium was exchanged for N2B27 medium (without CHIR99021) every 24h and gastruloids were grown up to 120h (5 days). Gastruloids were collected for sequencing at d3, d3.5, d4, d4.5 and d5. Gastruloids for half-day time points were set up in the evening to allow collection of full- and half-day time points at the same time.

### Sample preparation: scRNA-seq

Gastruloids were grown as described above. At d3 or d4, gastruloids were transferred to an eppendorf tube, washed with PBS and dissociated using Accutase (StemPro, A1110501) to obtain a single cell suspension. To remove Accutase, cells were washed twice in 5 ml PBS + 0.04% BSA (Gibco, 15260037) and filtered through a 50 um strainer (Sysmex, 1050553). Cell number and viability was assessed using the Countess II Automated Cell Counter and samples were diluted to submit for 10x 3’ scRNA sequencing at a targeted cell capture rate of 10,000 cells per lane.

### Sample preparation: MULTI-seq

Gastruloids were grown as described above. At the collection time point, up to 32 representative gastruloids were selected and individual gastruloids were transferred to wells of a U-bottom suspension culture 96-well plate with 80 μl Accutase (StemPro, A1110501) per well. Gastruloids from multiple time points were combined on a single plate to minimise batch effect. Gastruloids were incubated at 37 *°C* for three minutes and were dissociated through repeated pipetting using a multichannel pipette. Low-retention pipette tips were used for this and all subsequent steps to minimise cell loss. Incubation and pipetting were repeated until a near single-cell suspension was achieved in each well.

To label the cells of individual gastruloids with MULTI-seq barcodes, the MULTI-seq protocol (McGinnis et al., 2019) was adapted for low cell numbers as follows. First, 10 μl of unique 10x Anchor:Barcode solution was added to each well. Samples were mixed by pipetting and incubated on ice for 5 minutes. Henceforward, samples were kept on ice. Next, 10 μl 10x Co-Anchor solution was added, mixed in by pipetting and incubated on ice for 5 minutes. Anchor/Co-Anchor stock concentrations were calculated based on estimated cell numbers per gastruloid (7500 cells at d3, 13000 cells at d4, 25000 cells at d5). Used barcodes can be found in Supplementary table 1 (see supplementary note 1 for how these were selected). Barcode labelling was quenched by adding 100 μl ice cold PBS with 1% BSA (Gibco, 15260037) (PBS/BSA) per well. Next, cells from individual wells were combined into a 15 ml Falcon tube with 5 ml ice cold PBS/BSA to further quench barcode labelling. Cells were spun down at 300g in a precooled centrifuge at 4 °C. The cell pellet was washed with 5 ml ice cold PBS/BSA and spun down again. Next, the cell pellet was resuspended in 1 ml ice cold PBS/BSA and passed through a Flowmi cell strainer (Bel-Art, H13680-0040) into a clean precooled 15 ml Falcon tube. The volume was topped up to 5 ml with ice cold PBS/BSA for a final wash and the cells were spun down again. All but approximately 50 μl of PBS/BSA was aspirated and the cells were resuspended in the remaining volume. Cell number and viability was assessed using the Countess II Automated Cell Counter and samples were diluted to submit for 10x 3’ scRNA sequencing at a targeted cell capture rate of 10,000 cells per lane.

### Single cell MULTI-seq

After dissociation and labelling with MULTI-seq barcodes, cells were processed using 10x Genomics Chromium single cell 3’ RNA sequencing v3.1 according to manufacturer’s instructions. For MULTI-seq, the MULTI-seq primer was added during cDNA amplification and the supernatant was collected during 0.6x SPRI clean-up to obtain the barcode fraction as described by McGinnis *et al*. MULTI-seq barcode libraries were prepared according to McGinnis *et al*. Libraries were amplified between 11 and 16 times. The number of amplification cycles required was empirically determined for each sample. scRNA and barcode libraries were sequenced on either HiSeq or NovaSeq 6000 instruments according to 10x Genomics instructions (read 1, 28 cycles; index i7, 8 cycles; and read 2, 91 cycles).

Importantly, owing to MULTI-seq barcoding we were able to pool gastruloids from multiple time points across three experiments. The first experiment involved 24 day 5 gastruloids spread over 3 10X lanes. In the first experiment (“exp1_d5”), barcodes 1 to 24 were used. After the first experiment, the optimal barcodes were used, as selected using the method described above. The second experiment involved 3 pools of day 3 and day 3.5 gastruloids (“exp2A_d3_d3.5”, “exp2C_d3_d3.5”, “exp2D_d3_d3.5”; note pool B w as not sent for 10X sequencing), as well as 3 pools of day 4 and day 4.5 gastruloids (“exp3A_d4_d4.5”, “exp3B_d4_d4.5”, “exp3C_d4_d4.5”). Unfortunately, the day 3 and day 3.5 gastruloid data quality was too poor and these experiments were excluded. For the day 4 and day 4.5 gastruloids, out of the 3 pools (pool A, B, and C), only pools B and C were sequenced, and from these only lanes (samples) 2 and 3 from pool B, and lanes 1 and 2 from pool C. For the second experiment, the top 24 barcodes were used. The third experiment involved 1 pool of 24 day 3 gastruloids, spread across 2 10X lanes (“exp4_d3”), as well as one pool of 16 day 3.5 and 16 day 4 gastruloids, spread across 4 10X lanes (“exp5_d3.5_d4”), and finally 16 day 4.5 and 16 day 5 gastruloids, spread across 3 10X lanes (“exp6_d4.5_d5”). The MULTI-seq barcodes for this third experiment were additionally resequenced at higher depth and using 16 PCR amplification cycles during library preparation due to low yield.

### 10x Genomics data pre-processing

Raw FastQ files were processed with Cell Ranger 6.0.0 using default mapping arguments, and the mm10-2020-A genome and annotation (provided by 10X Genomics).

### Quality Control

CellRanger called cells were subjected to stringent quality control metrics on a sample-by-sample basis. They were selected for minimum number of UMI, minimum number of genes detected, as well as minimum and maximum mitochondrial percentage. Furthermore, cells that passed this initial QC were processed using the hybrid method from the scds doublet calling R package (Bais and Kostka, 2020). All cells that the method called as doublets were removed, along with any cell which had more than 10 of its 30 nearest neighbours called as doublets by the hybrid method. After stringent QC, we retained 77,683 cells. 72,176 of these cells were sequenced using MULTI-seq. The remaining 5,507 cells were sequenced using 3’ RNA sequencing.

For comparison purposes across experiments (shown in supplementary figure 1 B, D, F, and H), QC metrics were normalised across samples, to have equal mean and standard deviation. In the case of the mitochondrial percentage and the number of UMI this was done on the log2 of the values.

QC thresholds:

|  | 10X | MULTI-seq |  |  |
| --- | --- | --- | --- | --- |
|  | gastr_d3,<br>gastr_d4 | exp1 | exp3 | exp4,<br>exp5,<br>exp6 |
| Minimum number of UMI | 4000 |  | 8000 |  |
| Minimum number of genes | 1500 |  | 2500 |  |
| Minimum percent mitochondrial | 3 |  |  | 1 |
| Maximum percent mitochondrial | 10 |  |  |  |

### MULTI-seq demultiplexing

Cells were assigned MULTI-seq barcodes from the R1 and R3 fastq files using the published MULTI-seq demultiplexing method for barcode alignment and classification, with the slight modification that only cells whose barcodes passed RNA QC were included in the demultiplexing. This increased the sensitivity of the method. In all but the first MULTI-seq experiment the barcode yield was low, probably due to a loss of MULTI-seq anchor/co-anchor efficiency over time. However, it was still possible to classify these cells using the k-means based negative cell reclassification method as implemented in MULTI-seq for all but exp3C_d4_d4.5. A sample-specific reclassification stability was chosen based on identifying local maxima in diagnostic plots (no reclassification was done for exp1; for exp3B_d4_d4.5 sample2: 3; exp3B_d4_d4.5 sample3: 6; exp4_d3 sampleA: 6; exp4_d3 sampleB: 6; exp5_d3.5_d4 sampleA: 9; exp5_d3.5_d4 sampleA: 10; exp5_d3.5_d4 sampleA: 18; exp5_d3.5_d4 sampleA: 8; exp6_d4.5_d5 sampleA: 4; exp6_d4.5_d5 sampleB: 4; exp6_d4.5_d5 sampleC: 5). In experiment 4 many cells spuriously associated with barcode 6 and barcode 14 in sample A, and barcodes 12, 24, and 30 in sample B. All cells assigned to these barcodes whose top detected barcode was not the barcode they were assigned were reassigned as negative and included in the negative cell reclassification. The attractor barcodes were excluded from negative cell reclassification, as well as barcode 61 in sample A, and barcode 13 in sample B, as these also acted as spurious attractors during reclassification. These issues led to only around 50% of these cells being confidently assigned a barcode (Supplementary Figure S4C).

### Sample integration

Subsequently, the samples were concatenated, 1274 cell-cycle associated genes were removed based on their associated GO terms, and the samples normalised using multiBatchNorm from the batchelor R package (Haghverdi et al., 2018). Highly variable genes were selected using the modelGeneVar function from batchelor, with a minimum mean of 0.001 and a maximum p-value of 0.05. Subsequently, multiBatchPCA was used to calculate the first 50 PCs. The PCA space was then batch effect corrected using MNN, first correcting samples within a time point (ordered from most cells to fewest), and then correcting across time points (ordered from day 5 to day 3).

### Joint QC

The 50-dimensional corrected PCA space was clustered using Seurat Version 4’s FindNeighbors function with a k of 30, followed by the FindClusters function with a resolution of 1 (Hao et al., 2021). A low mitochondrial, high ribosomal cluster was subsequently further removed. The data was then reintegrated as above, but without cells contained in this low-quality cluster.

### Visualisation

A UMAP was generated using the “naive” method with default parameters from the R package umap version 0.2.8 (Konopka, Tomasz, 2022), based on the corrected 50-dimensional PCA space using random state 2402.

### Mapping to embryo atlas

The gastruloid data was integrated with the Pijuan-Sala (Pijuan-Sala et al., 2019) and the Imaz-Rosshandler extended (Imaz-Rosshandler et al., 2022) mouse gastrulation and organogenesis atlases. In both cases, and as previously described, cell cycle genes were removed for integration, and the two datasets were subset to common genes. For label transfer, each sample was mapped individually to the atlas; to obtain a joint embedding, all samples were mapped at the same time. The samples were normalised using multiBatchNorm from the batchelor R package version 1.6.3. Highly variable genes were selected using the modelGeneVar function from batchelor, with a minimum mean of 0.001 and a maximum p-value of 0.05. Subsequently multiBatchPCA from batchelor was used to calculate the first 50 PCs. The PCA space was then batch effect corrected using reducedMNN, first correcting the atlas (first samples within a time point from those with the most cells to those with the fewest cells, then across time points, in order of latest time point to earliest time point). In the case where all gastruloid samples were mapped at the same time, subsequently the gastruloid samples were iteratively mapped to the integrated PCA space, in order of last time point to first time point, and within each time point ordered based on number of cells (from most to fewest). In the case where the samples were mapped individually, the sample was integrated with the atlas PCA-space. Cell types were then assigned based on the majority vote of the 30 nearest atlas cells to each gastruloid cell. In the original atlas, these cell types were further refined using the Guibentif *et al*. somite reannotation (Guibentif et al., 2021). For this, the dataset was subset to all cells that were originally assigned as paraxial mesoderm, somitic mesoderm, or intermediate mesoderm, and remapped to just the annotated E8.5 atlas cells.

### Clustering and cell annotation

The initial reference-based cell type annotation was used, together with marker gene expression, to guide a *de novo* cell type annotation. For this, the gastruloid-only integrated PCA space was clustered using Seurat Version 4’s FindNeighbours function with a k of 30, and Seurat Version 4’s FindClusters function with a resolution of 0.6 and random seed of 0 (Hao et al., 2021). This yielded 16 clusters which were given preliminary annotations. Subsequently, the borders between cell types were refined, and clusters subclustered, in a biologically-informed, supervised manner. For this, each cell’s log-normalised gene expression was stabilised by averaging each gene over the cell’s 30 nearest neighbours.

The cluster annotated as primitive streak was further refined into PGCs, primitive streak, and epiblast, by subsetting the smoothed gene expression matrix to that cluster and the genes Dppa5a, Utf1, Zfp42, Pou5f1, Brachyury (T), and Wnt3. PCA dimensionality reduction was performed on this space, and k-means clustering using the first two principal components, and a k of 10 was performed, using 10 random initialisations. One of the resulting clusters was classified as PGCs, while a second was assigned epiblast. The remaining two clusters retained a primitive streak identity.

A similar procedure was done to distinguish between caudal neurectoderm, primitive streak, early neurectoderm, and caudal epiblast using Thy1, Hes3, Nrp2, Epha5, Gbx2, Sfrp1, Ncam1, Pou5f1, Hoxc8, Hoxb9, Hoxaas3, Pim2, T, and Wnt3 as marker genes. In this case a mixture of 4 ellipsoidal Gaussians of varying volume, shape, and orientation was fit, and each cluster assigned one cell type.

To distinguish between anterior primitive streak and nascent mesoderm, T, Foxa2, Cyp26a1, Lhx1, and Mesp1 were used as marker genes. Only the second principal component was used as an input to a k-means clustering with 10 random initialisations and a k of 2.

Similarly, to distinguish between nascent mesoderm and head mesoderm, Cyp26a1, Lhx1, Mesp1, Igfbp5, and Sox9 were used as marker genes, and the first two principal components used as input to fitting a mixture of 2 ellipsoidal Gaussians of varying volume, shape, and orientation.

The spinal cord was subset into early and late by reclustering the spinal cord cells using Seurat’s FindNeighbors (k=30), and FindClusters with a resolution of 0.2 and a random seed of 0.

Similarly we distinguished between early anterior PSM, late anterior PSM, and somites by clustering those cells using Seurat’s FindNeighbors (k=30) and FindClusters with a resolution of 0.3 and a random seed of 0.

We distinguished between Caudal Epiblast and NMPs using Pou5f1, Pim2, Cyp26a1, and Sox2 as marker genes, and the first two principal components used as input to fitting a mixture of 2 ellipsoidal Gaussians of varying volume, shape, and orientation. We further refined the distinction by requiring that all caudal epiblast cells have a stabilised Cyp26a1 expression of less than 0.1.

### Visualising Marker Gene Expression by Cell Type

To visualise marker gene expression, we used the normalised and 30-nn stabilised gene expression matrix as above. For each of the marker genes, we calculated the per cell type mean. We subsequently scaled each gene individually by subtracting the mean of the per cell type means, and dividing by the standard deviation of the per cell type means. This was then plotted as a heatmap.

### Calculating each cell’s equivalent embryonic day

For each gastruloid cell, the number of its 30-nearest cells of each embryonic day in the extended atlas (after individual sample integration) was recorded. This yielded a cell-by-embryonic day matrix, with each row summing to 30. This matrix was then normalised by the number of cells in the atlas that were of that embryonic day, to control for varying detection of each embryonic day. Thus the columns summed to 1. This was then divided by the rowSums, so that each gastruloid cell was equally weighted. Based on this, each gastruloid cell was assigned an embryonic day, calculated as the weighted average of the embryonic days (the sum of its row values multiplied by the column names).

### Comparison with Previous Gastruloid scRNA-seq Dataset

The van den Brink coutb UMI count tables were downloaded from GEO, along with the metadata. 458 cells that were in the metadata were not in any of the samples downloaded from GEO, from 2 CelSeq “bulk” samples. Furthermore, 7959 cells that were not in the metadata were in the count matrices, presumably cells that had been removed at the QC stage. All downloaded data were re-QCed using the above method, with custom, experiment-specific, thresholds. Based on visual inspection, it was clear that 4 samples had been assigned an incorrect batch, and were manually altered. Additionally, there were 2 batches with only 1 and 2 samples assigned respectively. Further investigation revealed that these were actually samples belonging to other, bigger batches. The QC-ed dataset was integrated with the present data, using the same code as had been used to integrate just the present data, except that the genes were subset to just common genes in the two datasets. A UMAP with random state 2402 was run on the first 30 PCs of the integrated PCA space. Colours were assigned to the van den Brink cell types to align with the closest match in the present data.

To quantitatively compare the similarity between the datasets, for each day 5 cell in the integrated dataset, the batch of its 30 nearest neighbours was recorded, to produce a cell-by-batch matrix with the rows summing to 30. The batches were the two day 5 experiments in the present data (sequenced using 10X), as well as the single van den Brink 10X experiment, and the IB10 and Lfng CEL-seq experiments. This was then divided by the number of cells in each experiment to control for the varying numbers of cells in each experiment, and then divided by the row sums so that each cell would be equally weighted. We then plotted this normalised nearest neighbour score.

### Trajectory Inference

Velocyto (La Manno et al., 2018) was run with default parameters for each chromosome separately, to obtain the spliced and unspliced count matrices. For trajectory inference, data were subset to only cells confidently assigned either a neural or a mesodermal gastruloid. The inference was performed separately for neural and mesodermal gastruloids, under the assumption that a cell in one gastruloid class will never give rise to a cell in another gastruloid class. All cell cycle-associated genes were removed.

First, the data were preprocessed according to scVelo (Bergen et al., 2020). The data were filtered and normalised using the top 2000 genes, with a minimum shared count of 20. Subsequently, the neighbourhood graph was constructed using the 50 PCs from the gastruloid-only integrated PCA space, and 30 nearest neighbours. This was also used to generate moments. scVelo was then run in the dynamical mode, and the velocity graph was approximated. CellRank (Lange et al., 2022) was used to turn the output into a kernel, and compute the transition matrix. For the flow plots, the resulting anndata objects were concatenated and their velocity embedding streams plotted. Figure 1G was produced manually.

The Waddington OT (Schiebinger et al., 2019) implementation in CellRank was used. For each cell, proliferation and apoptosis scores were calculated based on the marker genes provided in CellRank, and used to calculate initial growth rates. The Waddington OT transition matrices were then calculated using 10 growth iterations. The Waddington OT flow plots were generated from the resulting transition maps, separately for neural and mesodermal gastruloids. Additionally, the Waddington OT-estimated growth rates were separately plotted.

For the terminal state plots, CellRank was used to generate a connectivity kernel, using GPCCA, and a combination of 0.5 velocity, 0.4 Waddington, and 0.1 connectivity. Based on a Schur decomposition computing 20 macrostates, 10 macrostates were chosen for the neural gastruloids, and 12 for the mesodermal gastruloids. Terminal states were manually identified from macrostates, by excluding any macrostates not in the final time points (epiblast in the case of neural gastruloids, and one of the two mature endoderm macrostates in the case of the mesodermal gastruloids). Absorption probabilities were calculated using default parameters.

### PCA on gastruloid cell type proportions

To investigate the inter-gastruloid heterogeneity, we generated a count matrix of cell type by gastruloid, with entries corresponding to the number of cells of each cell type assigned to each gastruloid. We divided each row by the total number of cells in each gastruloid to obtain the proportion of cells in each gastruloid assigned a certain cell type. We then ran centred, but not scaled, PCA on this, to embed each gastruloid in a principal component space. We did this both for all gastruloids, as well as for each time point separately. When investigating the day 5 intermediate gastruloids, we used the same approach but subset to all day 4.5 and day 5 gastruloids.

Where gastruloids are listed in a linear order, they are sorted based on their principal curve in their PC space (the number of PCs used, between 2 and 4, was manually chosen based on elbow plots). This was calculated using the principal_curve function from R’s princurve package.

### Binning cell types into lineages

To simplify visualisations, cell types were given broader classifications as multipotent, mesoderm, endoderm, or ectoderm (Supplementary Figure 6Q): PGCs, Epiblast, Primitive Streak, Caudal Epiblast, and NMPs were classified as multipotent; Notochord, Early Nascent Mesoderm, Late Nascent Mesoderm, Head Mesoderm, Cardiopharyngeal Mesoderm, Endothelium, Early Posterior PSM, Late Posterior PSM, Early Anterior PSM, Late Anterior PSM, Somites, and Mature Somites were classified as mesodermal; Anterior Primitive Streak and Mature Endoderm were classified as endoderm; Caudal Neurectoderm, Early Spinal Cord, Late Spinal Cord, Early Neurectoderm, Late Neurectoderm, and Neurons were classified as ectoderm.

### AICc calculation

To statistically confirm the existence of multiple gastruloid classes, we calculated the corrected AIC of a binomial model based on a k-means clustering, with varying k. We used the corrected AIC to account for our small sample size, and a simplification into a binomial model to ensure that the number of gastruloids remained greater than the number of parameters to be fit. For this, we obtained the number of cells in each gastruloid that were mesodermal, and the number of cells in each gastruloid that were neural, based on the coarse cell type binning described above. At each time point, we classified the cells into n classes using k-means clustering with five random initialisations. We then calculated the binomial log likelihood for each of the n classes as follows: 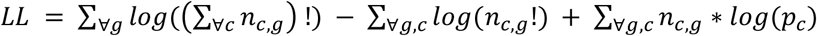, ∀_*g*_ assigned class *i* by the k-means clustering 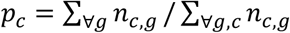 is the probability of each cell type (mesodermal or neural) in each class *n_c,g_* is the number of cells of a given cell type, *c*, that gastruloid *g* has, given that gastruloid *g* was assigned class *i*.

The model log likelihood was calculated as the sum of the individual class log likelihoods, and used as input to the AICc, with k=2*n classes: 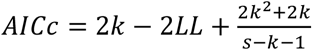, where *s* is the number of gastruloids.

### Separation into gastruloid classes

Based on the results of the AICc analysis, we manually classified the day 3 to day 4.5 gastruloids as either neural or mesodermal, and the day 5 gastruloids as neural, mesodermal, or intermediate. This was done based on visual inspection of the cell type proportion PCA plots, and agreed with the k–means clustering.

### Differential abundance testing between gastruloid classes

For each cell type at each time point, to compare if cell types are differentially abundant between two classes of gastruloid, we compared, for each cell type of interest, the proportion of cells in each gastruloid between each class using a Mann-Whitney U test. The p-values were then corrected for multiple hypothesis testing using a Benamini-Hochberg correction, and only comparisons with p<0.05 and FDR<0.1 displayed.

### scWGCNA

The day 3 gastruloid data were pseudobulked and cell cycle genes removed. The pseudobulk data was subsequently normalised to counts per million, and outlier genes removed using WGCNA’s goodSamplesGenes function (Langfelder and Horvath, 2008). The day 3 single cell data was then normalised, variable features selected, 50 PCs calculated, a neighbourhood graph constructed, and clusters found with a resolution of 1 using Seurat with default parameters. Subsequently, we followed the Feregrino and Tschopp method for scWGCNA (Feregrino and Tschopp, 2021).

### Significant module selection

The day 3 gastruloid data were randomly pseudobulked by permuting the gastruloid labels 10000 times. The amount of variance explained by each WGCNA-identified module eigengene in the pseudobulked gastruloid by gene matrix was then compared with the amount of variance that eigengene explained in the 10000 random pseudobulked matrices. This was done by using the variance explained in each of the 10000 random matrices as an empirical null distribution, and obtaining the p-value for the variance explained in the real data matrix. Modules that were significant at a Bonferoni-corrected 0.05 level were considered as explaining significant amounts of inter-gastruloid variance.

### GOEA

The significant modules were manually classified as late mesodermal (6 modules), early mesodermal (3 modules), late neural (6 modules), early neural (3 modules), and other (2 modules) based on their expression in the day 3 gastruloids. The 3 early neural modules were merged, and the 3 early mesodermal modules were merged. GO enrichment analysis was run separately on the mesodermal genes and the neural genes using enrichR (Xie et al., 2021) and the 2021 biological process annotation. Terms that were significant in either the mesodermal or the neural genes at an adjusted p-value of 0.05 (p-value adjustment performed in enrichR) were then compared using a Fisher exact test. In the Fisher exact test, the number of mesodermal genes in the GO term relative to the number of mesodermal genes was compared to the number of neural genes in the GO term relative to the number of neural genes. The resulting p-values were Benjamini-Hochberg corrected, and the terms that were significant at an FDR of 0.1 and a p-value of 0.05 reported.

### Comparison to bulk 2D differentiation data

Fastq files for the Gouti *et al*. 2014 paper were downloaded from the EBI Array express database (Gouti et al., 2014). Each sample was aligned to the mm10 genome using tophat followed by htseq. The count tables were then merged. The resulting count table was concatenated with the day 3 and day 3.5 gastruloid pseudobulk count matrices. These were then processed using DESeq, with the design matrix including only experiment of origin (with all Gouti *et al*. samples considered as coming from one experiment, and the MULTI-seq data separated based on whether they came from exp4_d3 or exp5_d3.5_d4) as a variable. Size factors were estimated, and genes whose total counts were less than 10 were removed. VST normalisation was performed, followed by Limma batch effect correction, with only organoid of origin (gastruloid or 2D) input as the batch variable. The resulting PCA space was plotted.

### Comparing day 4.5 and day 5 PCA distances

PCA was run on the day 4.5 and day 5 cell type proportion by gastruloid matrix as described above and the Euclidean distances between all gastruloids were calculated. For each day 5 gastruloid, its mean Euclidean distance to the day 4.5 mesodermal gastruloids was calculated, as well as its mean Euclidean distance to the day 4.5 neural gastruloids.

### Simulating perturbation experiments

We simulated perturbation experiments with varying numbers of samples per condition, and varying numbers of gastruloid per sample. For each gastruloid, the number of cells per gastruloid in each sample is generated from a Gamma-Poisson with a mean of 5000/’number of gastruloids to sample’ and an index of dispersion (IoD) of 500. Additionally, for each organoid the proportion of each cell type is obtained by sampling with replacement (i.e. bootstrapping) from the day 4.5 gastruloid cell type proportions. In the case where there is a fold change, this probability is additionally multiplied by the fold change and the resulting vector is renormalised to sum to 1 again. This probability vector is then multiplied by the previously obtained number of cells for that gastruloid. This was repeated for however many gastruloids there were in each sample, and again for however many samples were present in each experiment. For the simulations, 100 trials were run for each parameter combination.

### Poisson differential abundance testing

For Poisson differential abundance testing, the code was based on benchmarking code from miloR (Dann et al., 2022). A Poisson glm was fit for each cell type, with the covariates condition and offset. The offset was the log of the number of cells in that sample. Resulting p-values were Benjamini-Hochberg corrected, and all cell types whose FDR was less than 0.2 were classified as differentially abundant.

### Negative binomial differential abundance testing

For negative binomial differential abundance testing, again the code was based on benchmarking code from miloR (Dann et al., 2022). EdgeR normalisation factors were calculated using the TMM method, and subsequently edgeR was used to estimate dispersion for each cell type, and a quasi-likelihood model fit, and used for differential abundance testing. Again, all cell types whose FDR was less than 0.2 were classified as differentially abundant.

### Deconvolution differential abundance testing

The differential abundance testing approach was based on Young *et al*. code initially written for cell type deconvolution of bulk RNA-seq samples based on single cell references (Young et al., 2021). In this approach, prior knowledge from individual gastruloid sequencing is used to construct a PCA space for each gastruloid class, which is subsetted based on manual inspection of elbow plots. The PCA space is based on the log(x) of the cell type proportions, where any zero values of x are replaced by the minimum non-zero value of x, to allow for exponentiation of the fitted values to force the counts to be strictly positive. This accounts for the correlated, non-uniform variance structure.

For testing each cell type, that cell type was removed from the data, and a deconvolution model fit to the data. This deconvolution model modelled each sample’s cell type count vector as follows: 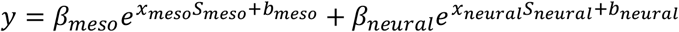, where the *β* scalars and *x* vectors are fit, and the S matrices and *b* vectors are the previously-calculated principal components and offsets of the respective mesodermal and neural gastruloid PCA spaces. *β* represents the number of cells that come from each respective gastruloid type, while *x* is the embedding of the sample in the respective PCA spaces S (this represents the mean of the pool of gastruloids of that class).

The model is fit by optimising a Poisson log-likelihood using stochastic gradient descent (SGD), adapting code from (Young et al., 2021). In particular, Adam, as implemented in tensorflow, was used for optimisation. As in the original implementation, the *β* scalars are fit as an exponential (*β* = *e^z^*) to enforce non-negativity, and two conditions are required for the optimisation to be terminated: The fractional decrease in the log likelihood needs to be less than some threshold parameter, and the fractional change in the sum of the sigmoid transformation of must be less than some sparsity threshold parameter. This second condition forces small *β* parameters to be optimised down to 0. The model is very resilient to initialisations. However, the *z* scalars were randomly initialised to the log of two positive values that sum to the total number of cells in the sample. The *x* vectors are intialised to 0.

After fitting the model, for each sample the error when applying the model for the tested cell type is recorded and a t-test performed on the residuals for the control versus the perturbed samples. The resulting p-values are then Benjamini-Hochberg corrected and all cell types whose adjusted p-value is less than 0.2 are classified as differentially abundant.

---

**Supplementary Figure 1 (related to Figure 1).**
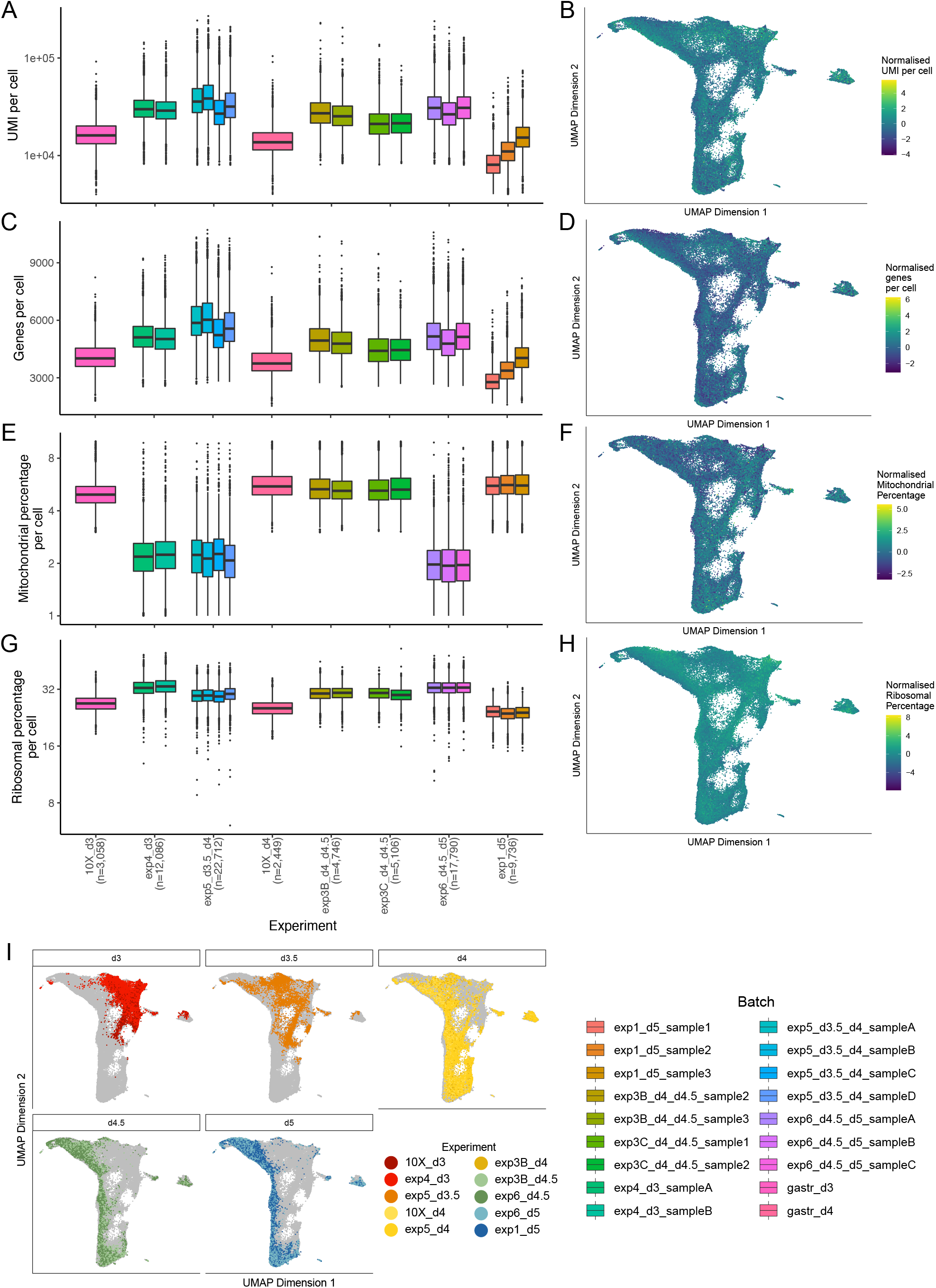
Quality Control Metrics for the scRNA-seq data. A - Box plot showing the number of unique molecular identifiers (UMI) per cell for each sample. Y axis is on a log2 scale. Samples are coloured based on the legend below. B - UMAP coloured by the log2 of the number of UMI per cell, normalised so each sample has the same mean and variance. C - Box plot showing the number of features (genes) per cell for each sample. Samples are coloured based on the below legend. D - UMAP coloured by the number of features (genes) per cell, normalised so each sample has the same mean and variance. E - Box plot showing the percentage of reads coming from mitochondrial genes per cell for each sample. Y axis is on a log2 scale. Samples are coloured based on the below legend. F - UMAP coloured by the log2 of the percentage of reads coming from mitochondrial genes per cell, normalised so each sample has the same mean and variance. G - Box plot showing the percentage of reads coming from ribosomal genes per cell for each sample. Y axis is on a log2 scale. Samples are coloured based on the below legend. H - UMAP coloured by percentage of reads coming from ribosomal genes per cell, normalised so each sample has the same mean and variance. I - UMAP showing all cells, with cells from specific time points highlighted, and coloured based on the experiment that they came from.

**Supplementary Figure 2 (related to Figure 1).**
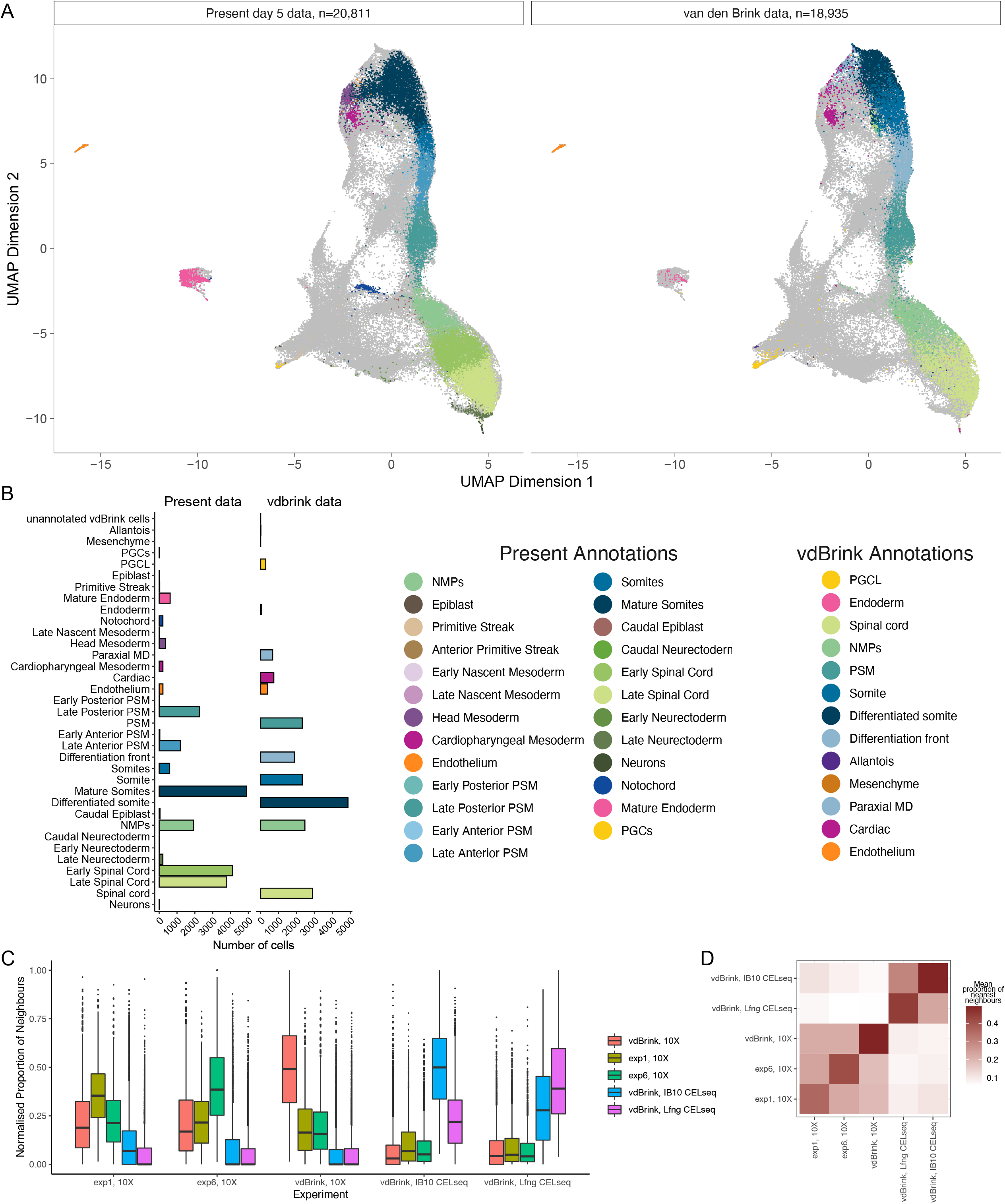
Comparison to previous scRNA-seq dataset of day 5 mouse gastruloids. A - UMAP of all cells from both datasets, with day 5 cells from each dataset highlighted, and coloured based on their respective time points. The separate cell type labels from both datasets were maintained but colours were chosen to highlight the parallels between cell types. B - Bar chart showing the number of day 5 gastruloid cells assigned each cell type. As above, cell type labels were chosen to ease comparison between datasets. C - Box plot showing the proportion of each cell’s neighbours (k=30) that come from a given experiment, after normalising for varying number of cells in each experiment. The x axis groups the data based on the experiment that the reference cell belonged to, and the data are coloured based on which experiment is being compared to. D - Heatmap showing the mean proportion of each cell’s neighbours that were from a given experiment. The reference cell’s experiment is on the x axis, and the experiment being compared to is on the y axis.

**Supplementary Figure 3 (related to Figure 1).**
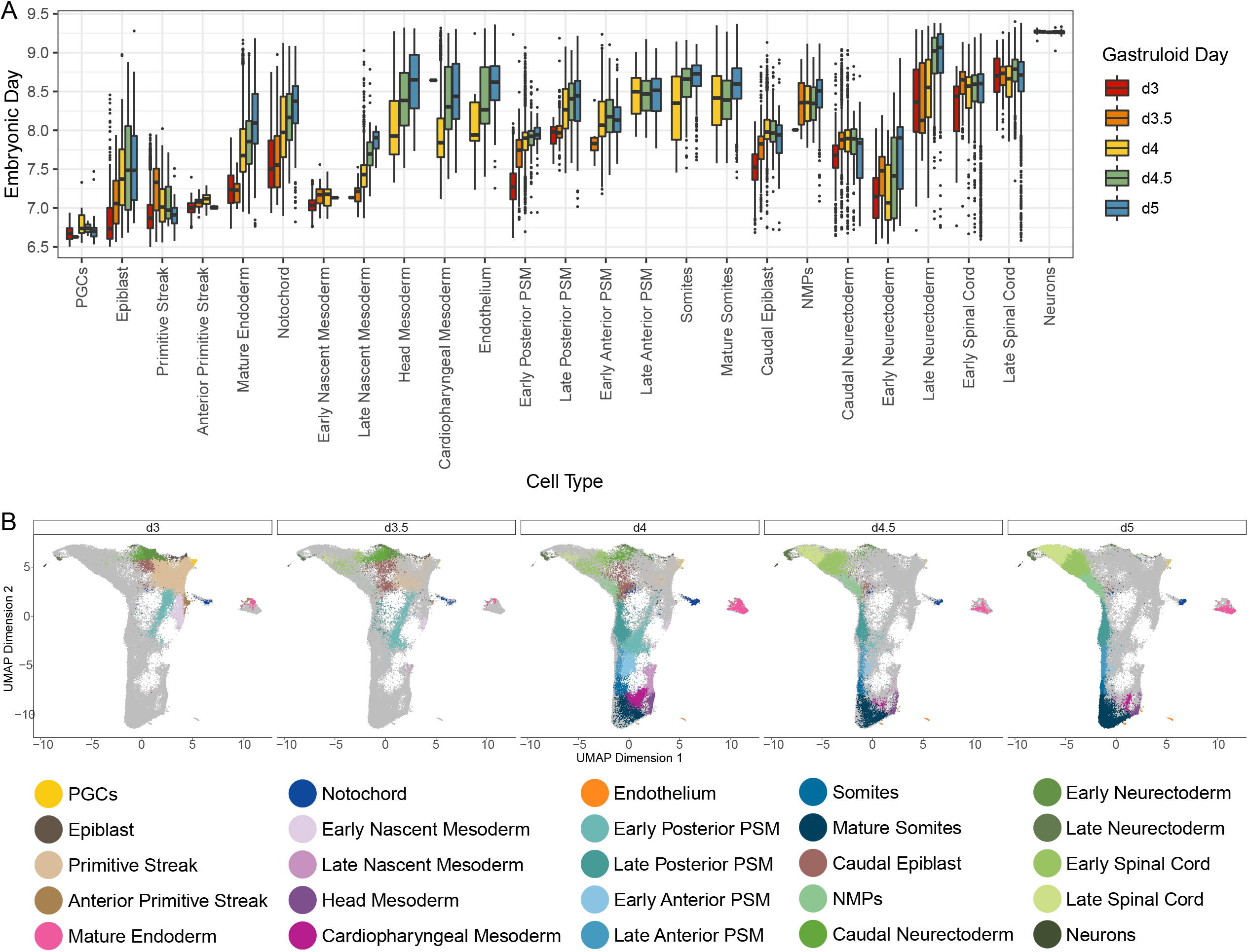
Different gastruloid cell types develop at different rates. A - Boxplot showing the embryonic day each gastruloid cell corresponds to, grouped by gastruloid sampling time point as well as gastruloid cell type B - UMAP of all cells, with cells from each time point highlighted separately, and coloured based on assigned cell type, according to the legend below.

**Supplementary Figure 4 (related to Figure 2).**
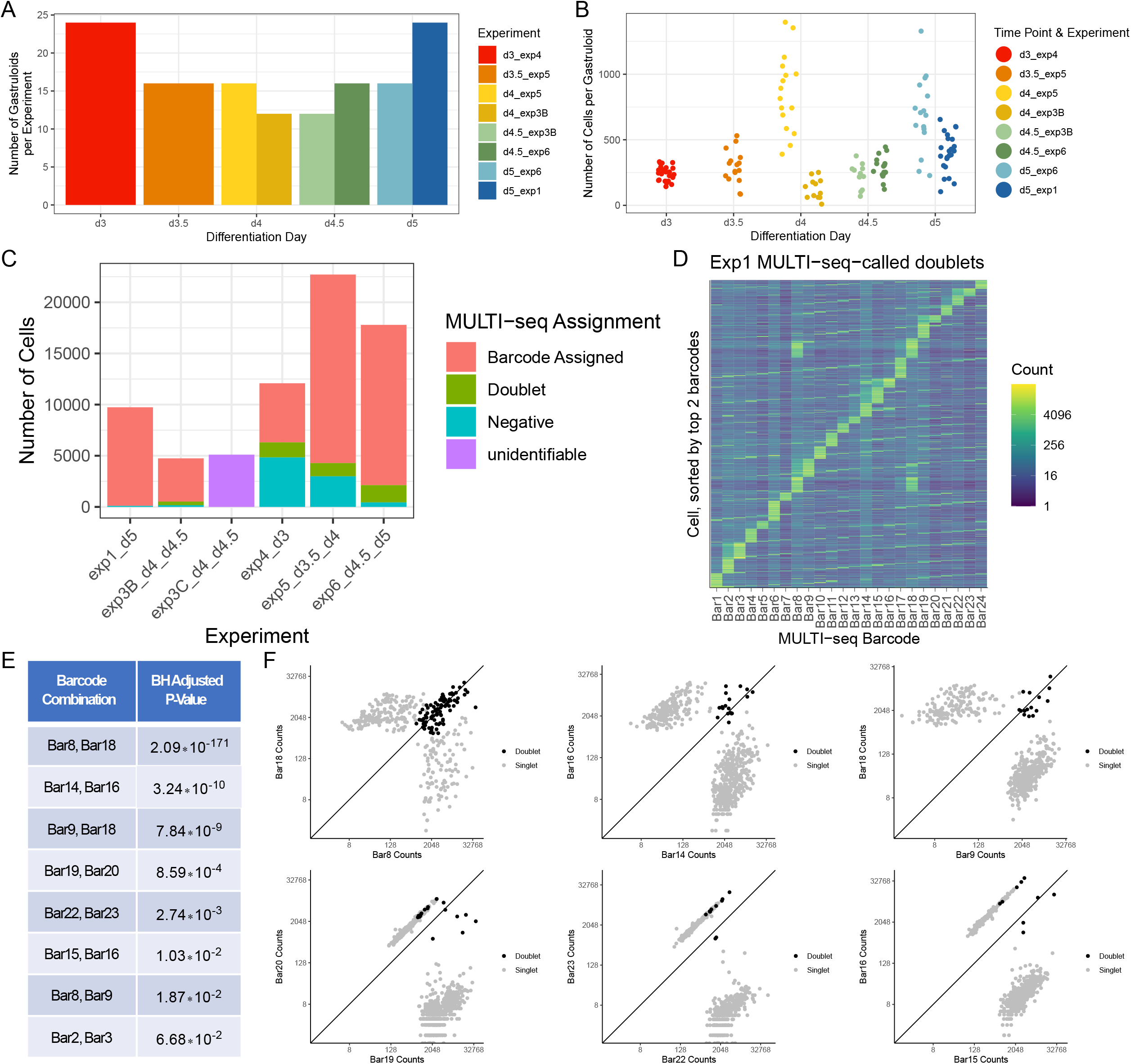
Quality Control Metrics for the MULTI- seq Data. A - Bar plot showing the number of gastruloids in each MULTI-seq experiment and time point (note that 3/5 experiments spanned two time points) B - Jitter plot showing the number of cells recovered for each gastruloid after MULTI-seq demultiplexing. The x axis shows the gastruloid’s time point, and the colour shows the time point and experiment of the gastruloid. C - Stacked box plot showing the number of cells from each experiment given each MULTI- seq assignment: A single unique barcode assigned, a MULTI-seq-called doublet (2 or more barcodes detected beyond their threshold values), a MULTI-seq called negative cell (no barcode detected above its threshold value), or unassigned, in the case of exp3C, for which the barcode data quality was not sufficient to attempt demultiplexing. D - A heatmap of the barcode expression counts in all the cells in exp1 that MULTI-seq identified as doublets, sorted on the y axis based on each cell’s top 2 barcodes. E - Table with the p-value of all barcode pairs that were overrepresented in the MULTI-seq- called doublets, based on an FDR threshold of 0.1. F - Plot comparing the counts of six of the overrepresented barcode pairs for singlets of each barcode, and doublets of both barcodes.

**Supplementary Figure 5 (related to Figure 2).**
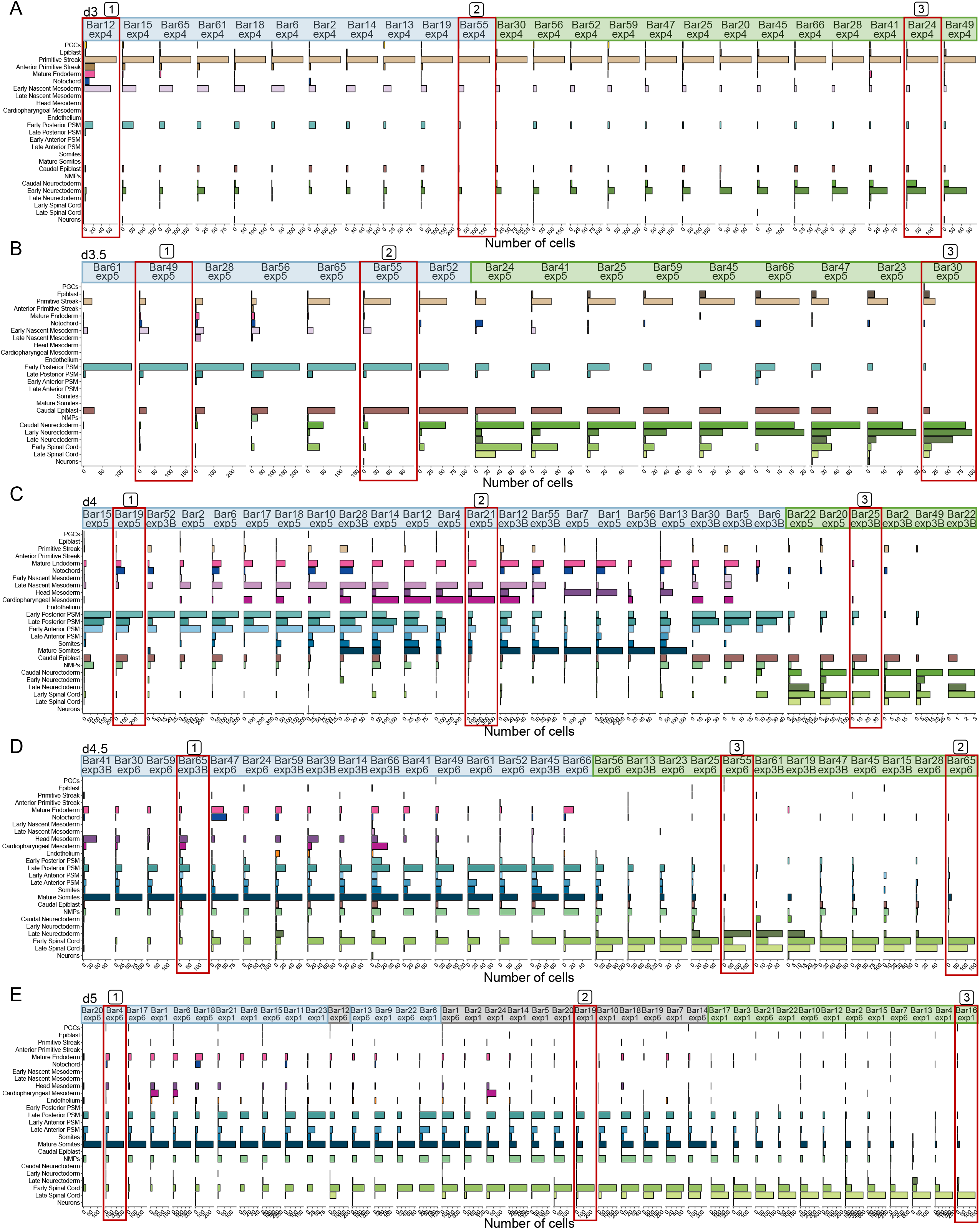
Bar plots of cell type abundance for each gastruloid. A-E - Bar plot showing the number of cells assigned to each cell type in each individual gastruloid, split by time point. Gastruloids are coloured based on whether they were assigned as mesodermal, neural, or intermediate, and the gastruloids selected as representative in 2D are highlighted. Gastruloids were sorted based on their ordering along the principal curve in their time point’s cell type proportion PCA space. Representative gastruloids shown in Fig. 2D are highlighted with red boxes and labelled.

**Supplementary Figure 6(related to Figure 2).**
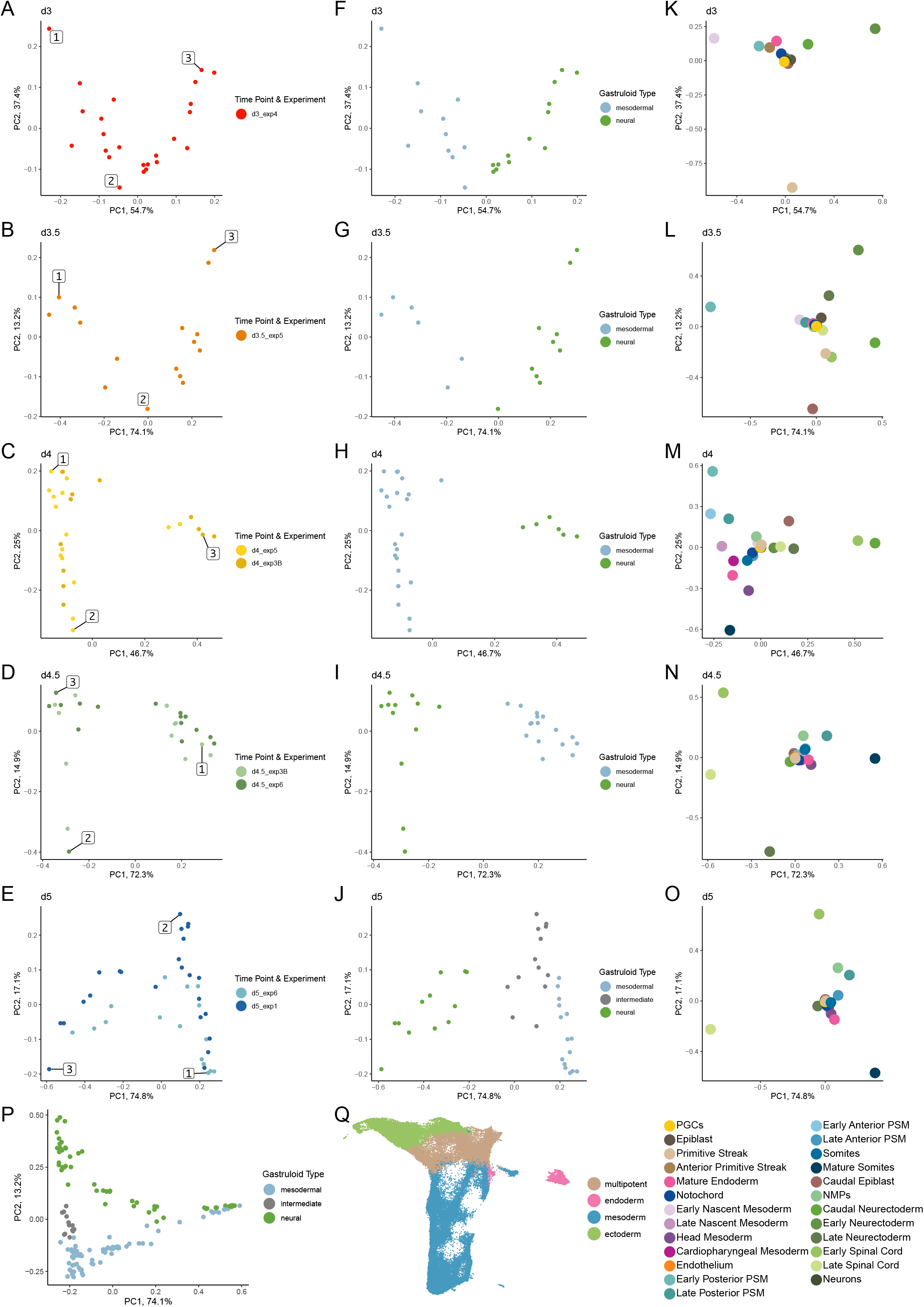
PCA space for each gastruloid. A-E - Embeddings of individual gastruloids from a given time point in a PCA space calculated based on the proportion of each gastruloid’s cells assigned each cell type. Gastruloids are coloured based on the time point and experiment that they came from. Representative gastruloids shown in Fig. 2D are labelled. F-J - Same embeddings as A-E but coloured by the class assigned to each gastruloid K-O - Feature loadings of the PCA spaces shown in (A-J). Each point is a cell type, coloured according to the legend below. P - Full PCA space based on all time points, as in Fig. 2B, coloured based on the class assigned to each gastruloid Q - UMAP of all cells showing the lineages each cell is assigned

**Supplementary Figure 7 (related to Figure 2).**
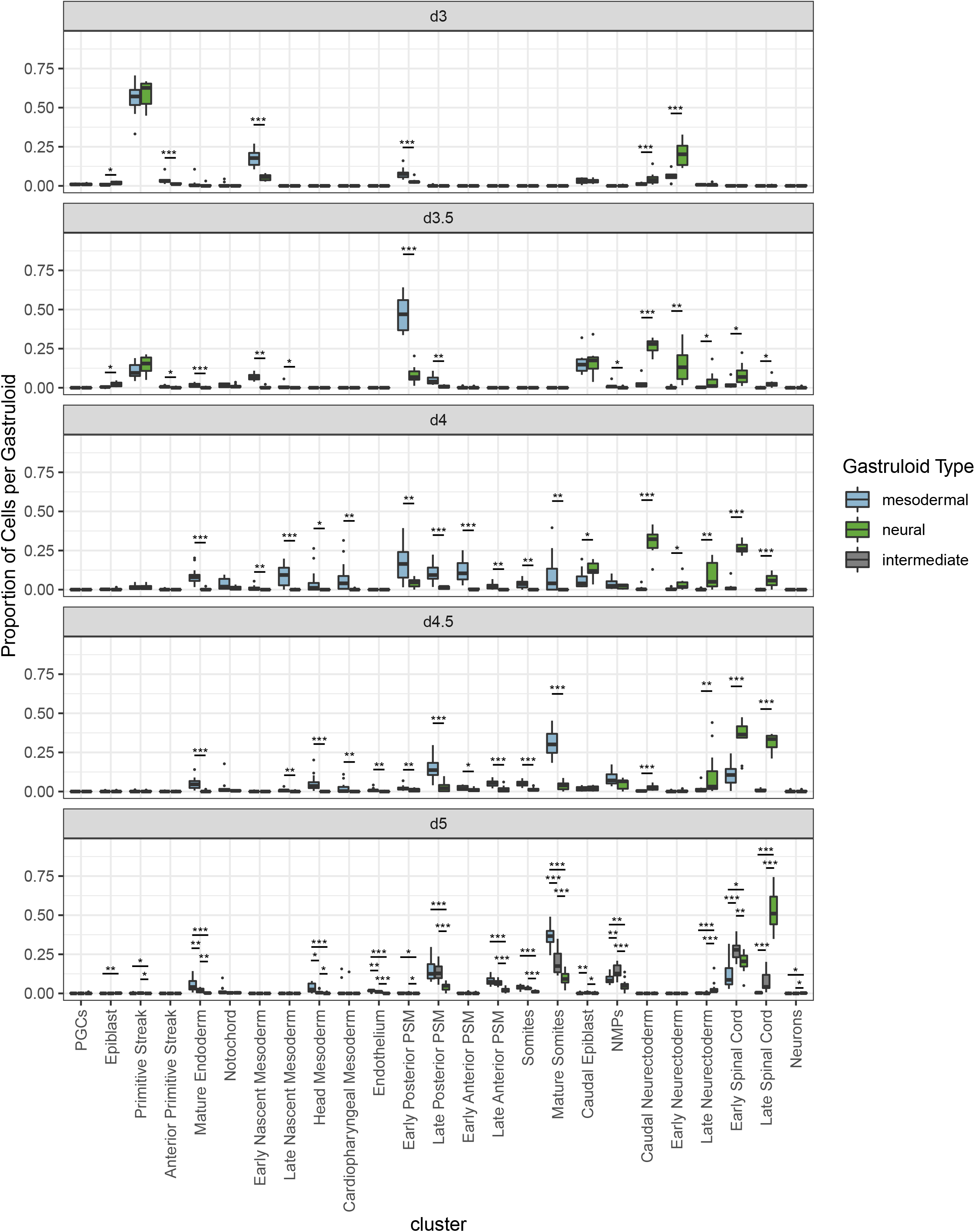
Differential abundance testing between gastruloid classes. Box plot showing proportion of cells of a given cell type in each gastruloid. Boxes are split by gastruloid time point and cell type, and coloured by gastruloid class. Stars indicate significance levels of comparisons between gastruloid classes based on a Wilcox rank sum test (***: p=0- 0.001; **: p=0.001-0.01; *: p=0.01-0.05), only comparisons with an FDR<0.1 are annotated.

**Supplementary Figure 8 (related to Figure 2).**
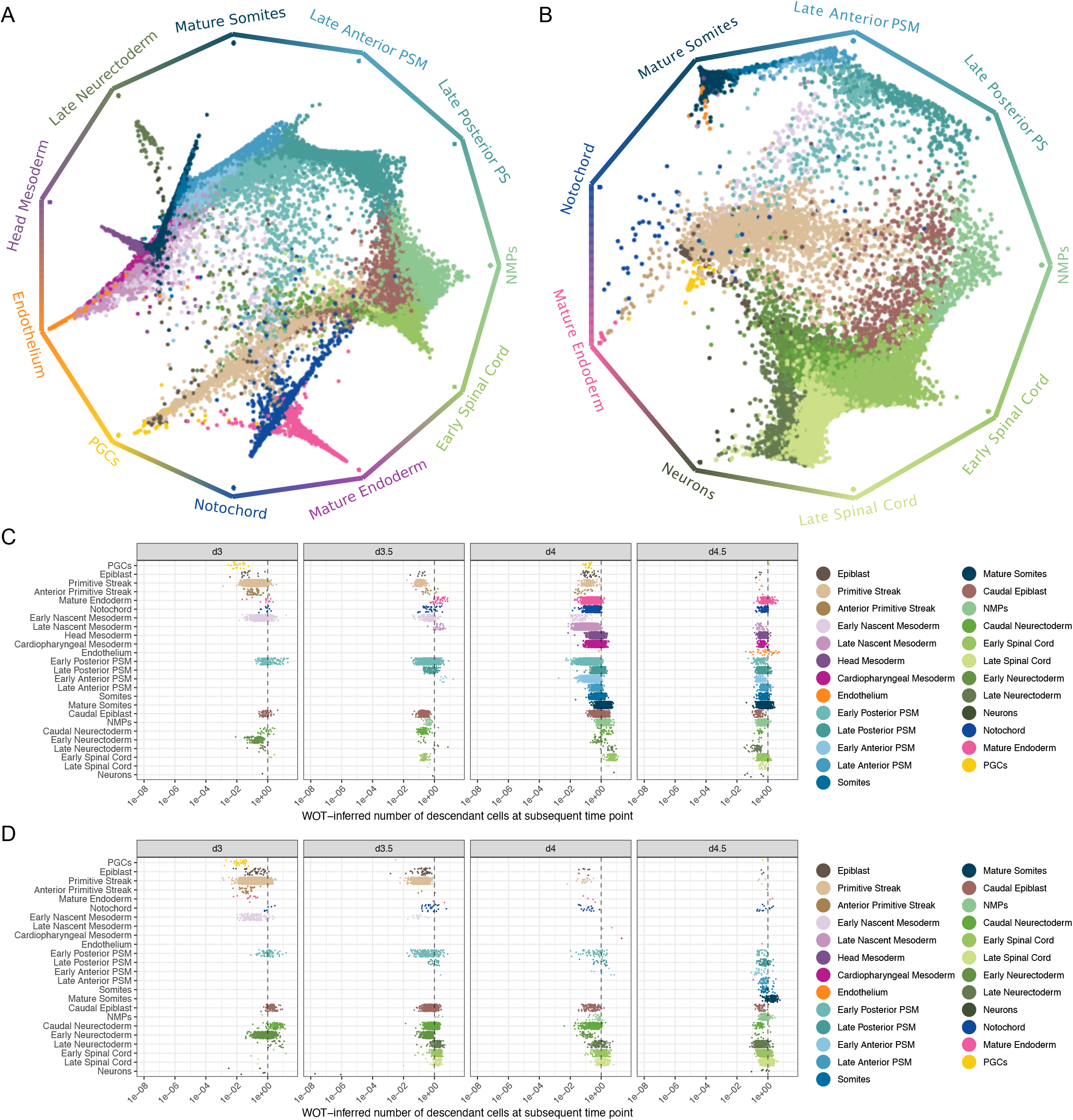
Separate trajectory inference for each gastruloid class. A - jitter plot showing the square root of the WOT-inferred estimated growth rates for each cell assigned a mesodermal gastruloid (roughly equivalent to estimated number of descendant cells), grouped and coloured based on assigned cell type, and split based on time point. Dotted line indicates 1 descendant cell. B - As A, but for cells assigned a neural gastruloid. C - CellRank terminal state plots showing the probability of each cell assigned a mesodermal gastruloid to be absorbed by a given terminal stage. B - As C, but for cells assigned a neural gastruloid.

**Supplementary Figure 9 (related to Figure 4).**
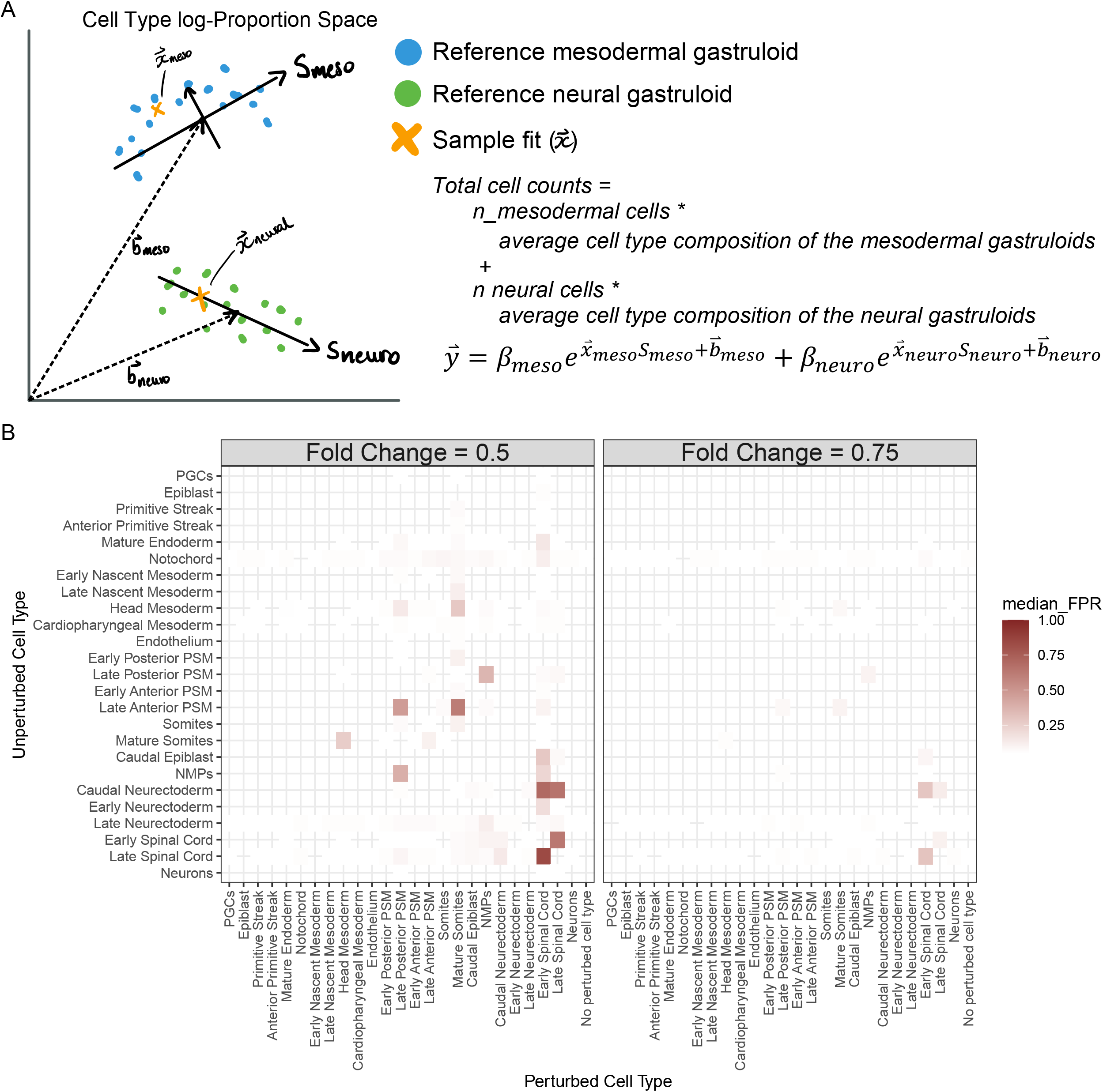
Systematic false positives in the deconvolution-based approach. A - Schematic of the deconvolution-based differential abundance testing method. Counts are modelled as the linear combination of gastruloids from the two classes. Within each class, the null distribution of gastruloids is modelled as falling along a PCA-space continuum (*S*), with offset (*β*) and some number of dimensions. The sample embedding location is represented as *x*. B - The deconvolution-based approach leads to systematic false positives when one cell type is perturbed owing to the correlation between cell types. The heatmaps show the median false positive rate across varying numbers of samples and gastruloids, for each cell type on the y axis, when a given cell type is perturbed (on the x axis). The data are shown for the x-axis cell type being perturbed at a fold change of 0.5 and at a fold change of 0.75. Median FPRs below 0.05 are removed. Each false positive rate is based on 100 simulations.

## Supplementary Notes

### Supplementary Note 1 - MULTI-seq Barcode Selection

In the first MULTI-seq experiment (exp1_d5), the first 24 MULTI-seq barcodes from McGinnis *et al*. were used. However, after demultiplexing, we observed that a large number of MULTI- seq doublets were detected (Supplementary Figure S5D). We found that eight pairs of barcodes were overrepresented according to the following hypothesis test, after Benjamini- Hochberg correction, and at an FDR threshold of 0.1 (Supplementary Figure S4E)

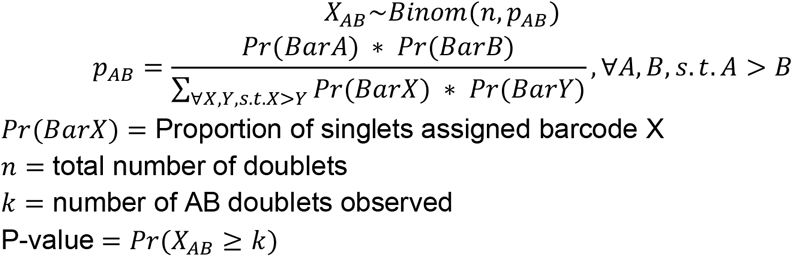

Barcodes 8 and 18 were most overrepresented. Inspection of their sequence revealed that they were reverse complements of each other. Additionally, barcodes 14 and 16 were also overrepresented, and they were 2 basepairs off from being reverse complements of each other. No other barcode pair was 2 or fewer base pairs from being reverse complements. The remaining overrepresented barcode pairs were sequential, aside from barcodes 9 and 18.

To explore this further, we plotted, for each barcode pair AB, the raw barcode counts for the barcode A singlets, barcode B singlets, and barcode AB MULTI-seq-called doublets (Supplementary Figure S4F). This revealed that the 8 and 18 MULTI-seq-called doublets, as well as the 14 and 16, and 9 and 18 MULTI-seq called doublets seemed to be true doublets, while the sequential MULTI-seq called doublets did not seem to be true doublets, but rather that there was contamination of the previous barcode in the subsequent barcode. The only plausible hypothesis for these two behaviours is that the reverse complement barcodes cause cells to stick together, forming true doublets, while when pooling the gastruloids, the previous barcode was not fully quenched, thus leading to contamination. This explains the behaviour of barcode 9 and barcode 18 MULTI-seq-called doublets: Contamination of barcode 8 in cells from gastruloid 9 caused them to anneal to cells from gastruloid 18, thus forming true doublets. As a result, in subsequent experiments we used barcodes selected to be at least 4 base pairs off of being reverse complements (see above), and made sure to use new pipette tips for each individual sample when pooling.

To avoid this issue, barcodes were selected to be optimally distant. Starting from the set of possible MULTI-seq barcodes identified by McGinnins *et al*., Hamming, Levenshtein, and longest common substring distances were calculated between barcodes; additionally the same distances were computed between barcodes and the reverse complement of all other barcodes. These were then given the following individual penalty functions:

- Hamming, Levenshtein, and reverse complement Levenshtein distance (given the least weight):

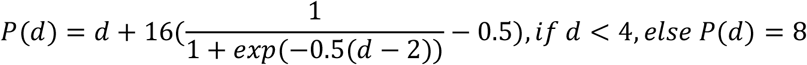
- Reverse complement hamming distance:

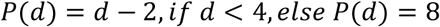
- Substring distance (weighted as important):

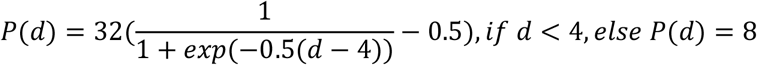
- Reverse complement substring distance (weighted as most important):

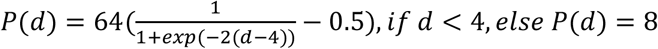

The minimum distance after penalty function application was used in a barcodebarcode distance matrix. Barcodes were then selected using a greedy algorithm: At each iteration, the barcode with the lowest sum score was removed. From this, the barcodes that were eliminated last were selected. However, owing to a previous computational error, the 32 optimal barcodes used were selected from a smaller set of barcodes that had already been purchased (1, 2, 3, 4, 5, 6, 7, 8, 9, 10, 11, 12, 13, 14, 15, 16, 17, 18, 19, 20, 21, 22, 23, 24, 25, 28, 30, 39, 41, 45, 47, 49, 52, 55, 56, 59, 61, 65, 66). These 32 barcodes were used for all subsequent MULTI-seq experiments.

## References

Anlaş, K., Gritti, N., Oriola, D., Arató, K., Nakaki, F., Lim, J.L., Sharpe, J., and Trivedi, V. (2021). Dynamics of anteroposterior axis establishment in a mammalian embryo-like system (Developmental Biology).

Argelaguet, R., Clark, S.J., Mohammed, H., Stapel, L.C., Krueger, C., Kapourani, C.-A., Imaz-Rosshandler, I., Lohoff, T., Xiang, Y., Hanna, C.W., et al. (2019). Multi-omics profiling of mouse gastrulation at single-cell resolution. Nature 576, 487–491. https://doi.org/10.1038/s41586-019-1825-8.

Baillie-Johnson, P., van den Brink, S.C., Balayo, T., Turner, D.A., and Martinez Arias, A. (2015). Generation of Aggregates of Mouse Embryonic Stem Cells that Show Symmetry Breaking, Polarization and Emergent Collective Behaviour In Vitro. J. Vis. Exp. 53252. https://doi.org/10.3791/53252.

Bais, A.S., and Kostka, D. (2020). scds: computational annotation of doublets in single-cell RNA sequencing data. Bioinformatics 36, 1150–1158. https://doi.org/10.1093/bioinformatics/btz698.

Barker, C.G., Petsalaki, E., Giudice, G., Sero, J., Ekpenyong, E.N., Bakal, C., and Petsalaki, E. (2022). Identification of phenotype-specific networks from paired gene expression–cell shape imaging data. Genome Res. 32, 750–765. https://doi.org/10.1101/gr.276059.121.

Beccari, L., Moris, N., Girgin, M., Turner, D.A., Baillie-Johnson, P., Cossy, A.-C., Lutolf, M.P., Duboule, D., and Arias, A.M. (2018). Multi-axial self-organization properties of mouse embryonic stem cells into gastruloids. Nature 562, 272–276. https://doi.org/10.1038/s41586-018-0578-0.

Bergen, V., Lange, M., Peidli, S., Wolf, F.A., and Theis, F.J. (2020). Generalizing RNA velocity to transient cell states through dynamical modeling. Nat. Biotechnol. 38, 1408–1414. https://doi.org/10.1038/s41587-020-0591-3.

van den Brink, S.C., Baillie-Johnson, P., Balayo, T., Hadjantonakis, A.-K., Nowotschin, S., Turner, D.A., and Martinez Arias, A. (2014). Symmetry breaking, germ layer specification *and axial organisation in aggregates of mouse embryonic stem cells*. Development 141, 4231–4242. https://doi.org/10.1242/dev.113001.

van den Brink, S.C., Alemany, A., van Batenburg, V., Moris, N., Blotenburg, M., Vivié, J., Baillie-Johnson, P., Nichols, J., Sonnen, K.F., Martinez Arias, A., et al. (2020). Single-cell and spatial transcriptomics reveal somitogenesis in gastruloids. Nature 582, 405–409. https://doi.org/10.1038/s41586-020-2024-3.

Criss, Z.K., Bhasin, N., Di Rienzi, S.C., Rajan, A., Deans-Fielder, K., Swaminathan, G., Kamyabi, N., Zeng, X.-L., Doddapaneni, H., Menon, V.K., et al. (2021). Drivers of transcriptional variance in human intestinal epithelial organoids. Physiol. Genomics 53, 486–508. https://doi.org/10.1152/physiolgenomics.00061.2021.

Dann, E., Henderson, N.C., Teichmann, S.A., Morgan, M.D., and Marioni, J.C. (2022). Differential abundance testing on single-cell data using k-nearest neighbor graphs. Nat.Biotechnol. 40, 245–253. https://doi.org/10.1038/s41587-021-01033-z.

Edri, S., Hayward, P., Baillie-Johnson, P., Steventon, B., and Arias, A.M. (2019). An Epiblast Stem Cell derived multipotent progenitor population for axial extension. Development dev. 168187. https://doi.org/10.1242/dev.168187.

Feregrino, C., and Tschopp, P. (2021). Assessing evolutionary and developmen taltranscriptome dynamics in homologous cell types. Dev. Dyn. dvdy. 384. https://doi.org/10.1002/dvdy.384.

Fleck, J.S., Sanchís-Calleja, F., He, Z., Santel, M., Boyle, M.J., Camp, J.G., and Treutlein, B. (2021). Resolving organoid brain region identities by mapping single-cell genomic data to reference atlases. Cell Stem Cell 28, 1148–1159.e8. https://doi.org/10.1016/j.stem.2021.02.015.

Gehling, K., Parekh, S., Schneider, F., Kirchner, M., Kondylis, V., Nikopoulou, C., and Tessarz, P. (2021). Single organoid RNA-sequencing reveals high organoid-to-organoid variability (Cell Biology).

Gouti, M., Tsakiridis, A., Wymeersch, F.J., Huang, Y., Kleinjung, J., Wilson, V., and Briscoe, J. (2014). In Vitro Generation of Neuromesodermal Progenitors Reveals Distinct Roles for Wnt Signalling in the Specification of Spinal Cord and Paraxial Mesoderm Identity. PLoS Biol. 12, e1001937. https://doi.org/10.1371/journal.pbio.1001937.

Gouti, M., Delile, J., Stamataki, D., Wymeersch, F.J., Huang, Y., Kleinjung, J., Wilson, V.,and Briscoe, J. (2017). A Gene Regulatory Network Balances Neural and Mesoderm Specification during Vertebrate Trunk Development. Dev. Cell 41, 243–261.e7. https://doi.org/10.1016/j.devcel.2017.04.002.

Guibentif, C., Griffiths, J.A., Imaz-Rosshandler, I., Ghazanfar, S., Nichols, J., Wilson, V., Göttgens, B., and Marioni, J.C. (2021). Diverse Routes toward Early Somites in the Mouse Embryo. Dev. Cell 56, 141–153.e6. https://doi.org/10.1016/j.devcel.2020.11.013.

Haber, A.L., Biton, M., Rogel, N., Herbst, R.H., Shekhar, K., Smillie, C., Burgin, G., Delorey, T.M., Howitt, M.R., Katz, Y., et al. (2017). A single-cell survey of the small intestinal epithelium. Nature 551, 333–339. https://doi.org/10.1038/nature24489.

Haghverdi, L., Lun, A.T.L., Morgan, M.D., and Marioni, J.C. (2018). Batch effects in singlecell RNA-sequencing data are corrected by matching mutual nearest neighbors. Nat.Biotechnol. 36, 421–427. https://doi.org/10.1038/nbt.4091.

Hao, Y., Hao, S., Andersen-Nissen, E., Mauck, W.M., Zheng, S., Butler, A., Lee, M.J., Wilk, A.J., Darby, C., Zager, M., et al. (2021). Integrated analysis of multimodal single-cell data. Cell 184, 3573–3587.e29. https://doi.org/10.1016/j.cell.2021.04.048.

Hof, L., Moreth, T., Koch, M., Liebisch, T., Kurtz, M., Tarnick, J., Lissek, S.M., Verstegen, M. M.A., van der Laan, L.J.W., Huch, M., et al. (2021). Long-term live imaging and multiscale analysis identify heterogeneity and core principles of epithelial organoid morphogenesis. BMC Biol. 19, 37. https://doi.org/10.1186/s12915-021-00958-w.

Imaz-Rosshandler, I., Rode, C., Guibentif, C., Ton, M.-L., Dhapola, P., Keitley, D., Argelaguet, R., Karlsson, G., de Brujin, M., Marioni, J., et al. (2022). Tracking Early Mammalian Organogenesis – Prediction and Validation of Differentiation Trajectories at Whole Organism Scale. Manuscr. Prep. https://marionilab.github.io/ExtendedMouseAtlas.

Konopka, Tomasz (2022). umap: Uniform Manifold Approximation and Projection.

La Manno, G., Soldatov, R., Zeisel, A., Braun, E., Hochgerner, H., Petukhov, V., Lidschreiber, K., Kastriti, M.E., Lönnerberg, P., Furlan, A., et al. (2018). RNA velocity of single cells. Nature 560, 494–498. https://doi.org/10.1038/s41586-018-0414-6.

Lange, M., Bergen, V., Klein, M., Setty, M., Reuter, B., Bakhti, M., Lickert, H., Ansari, M., Schniering, J., Schiller, H.B., et al. (2022). CellRank for directed single-cell fate mapping.Nat. Methods 19, 159–170. https://doi.org/10.1038/s41592-021-01346-6.

Langfelder, P., and Horvath, S. (2008). WGCNA: an R package for weighted correlation network analysis. BMC Bioinformatics 9, 559. https://doi.org/10.1186/1471-2105-9-559.

McGinnis, C.S., Patterson, D.M., Winkler, J., Conrad, D.N., Hein, M.Y., Srivastava, V., Hu, J.L., Murrow, L.M., Weissman, J.S., Werb, Z., et al. (2019). MULTI-seq: sample multiplexing for single-cell RNA sequencing using lipid-tagged indices. Nat. Methods 16, 619–626. https://doi.org/10.1038/s41592-019-0433-8.

Mittnenzweig, M., Mayshar, Y., Cheng, S., Ben-Yair, R., Hadas, R., Rais, Y., Chomsky, E., Reines, N., Uzonyi, A., Lumerman, L., et al. (2021). A single-embryo, single-cell time-resolved model for mouse gastrulation. Cell 184, 2825–2842.e22. https://doi.org/10.1016/j.cell.2021.04.004.

Mohammadi, S., Morell-Perez, C., Wright, C.W., Wyche, T.P., White, C.H., Sana, T.R., and Lieberman, L.A. (2021). Assessing donor-to-donor variability in human intestinal organoid cultures. Stem Cell Rep. 16, 2364–2378. https://doi.org/10.1016/j.stemcr.2021.07.016.

Pijuan-Sala, B., Griffiths, J.A., Guibentif, C., Hiscock, T.W., Jawaid, W., Calero-Nieto, F.J., Mulas, C., Ibarra-Soria, X., Tyser, R.C.V., Ho, D.L.L., et al. (2019). A single-cell molecular map of mouse gastrulation and early organogenesis. Nature 566, 490–495. https://doi.org/10.1038/s41586-019-0933-9.

Quadrato, G., Nguyen, T., Macosko, E.Z., Sherwood, J.L., Min Yang, S., Berger, D.R., Maria, N., Scholvin, J., Goldman, M., Kinney, J.P., et al. (2017). Cell diversity and network dynamics in photosensitive human brain organoids. Nature 545, 48–53. https://doi.org/10.1038/nature22047.

Ramachandran, P., Dobie, R., Wilson-Kanamori, J.R., Dora, E.F., Henderson, B.E.P., Luu, N. T., Portman, J.R., Matchett, K.P., Brice, M., Marwick, J.A., et al. (2019). Resolving the fibrotic niche of human liver cirrhosis at single-cell level. Nature 575, 512–518. https://doi.org/10.1038/s41586-019-1631-3.

Rossi, G., Broguiere, N., Miyamoto, M., Boni, A., Guiet, R., Girgin, M., Kelly, R.G., Kwon, C.,and Lutolf, M.P. (2021). Capturing Cardiogenesis in Gastruloids. Cell Stem Cell 28, 230–240.e6. https://doi.org/10.1016/j.stem.2020.10.013.

Sáez, M., Blassberg, R., Camacho-Aguilar, E., Siggia, E.D., Rand, D.A., and Briscoe, J. (2022). Statistically derived geometrical landscapes capture principles of decision-making dynamics during cell fate transitions. Cell Syst. 13, 12–28.e3. https://doi.org/10.1016/j.cels.2021.08.013.

Schiebinger, G., Shu, J., Tabaka, M., Cleary, B., Subramanian, V., Solomon, A., Gould, J., Liu, S., Lin, S., Berube, P., et al. (2019). Optimal-Transport Analysis of Single-Cell Gene Expression Identifies Developmental Trajectories in Reprogramming. Cell 176, 928–943.e22. https://doi.org/10.1016/j.cell.2019.01.006.

Tropepe, V., Hitoshi, S., Sirard, C., Mak, T.W., Rossant, J., and van der Kooy, D. (2001). Direct Neural Fate Specification from Embryonic Stem Cells. Neuron 30, 65–78. https://doi.org/10.1016/S0896-6273(01)00263-X.

Veenvliet, J.V., Bolondi, A., Kretzmer, H., Haut, L., Scholze-Wittler, M., Schifferl, D., Koch, F., Guignard, L., Kumar, A.S., Pustet, M., et al. (2020). Mouse embryonic stem cells selforganize into trunk-like structures with neural tube and somites. Science 370, eaba4937. https://doi.org/10.1126/science.aba4937.

Velasco, S., Kedaigle, A.J., Simmons, S.K., Nash, A., Rocha, M., Quadrato, G., Paulsen, B., Nguyen, L., Adiconis, X., Regev, A., et al. (2019). Individual brain organoids reproducibly form cell diversity of the human cerebral cortex. Nature 570, 523–527. https://doi.org/10.1038/s41586-019-1289-x.

Xie, Z., Bailey, A., Kuleshov, M.V., Clarke, D.J.B., Evangelista, J.E., Jenkins, S.L., Lachmann, A., Wojciechowicz, M.L., Kropiwnicki, E., Jagodnik, K.M., et al. (2021). Gene Set Knowledge Discovery with Enrichr. Curr. Protoc. 1. https://doi.org/10.1002/cpz1.90.

Young, M.D., Mitchell, T.J., Custers, L., Margaritis, T., Morales-Rodriguez, F., Kwakwa, K., Khabirova, E., Kildisiute, G., Oliver, T.R.W., de Krijger, R.R., et al. (2021). Single cell derived mRNA signals across human kidney tumors. Nat. Commun. 12, 3896. https://doi.org/10.1038/s41467-021-23949-5.

